# *Shigella* induces epigenetic reprogramming of zebrafish neutrophils

**DOI:** 10.1101/2022.11.03.515069

**Authors:** Margarida C. Gomes, Dominik Brokatzky, Magdalena K. Bielecka, Fiona C. Wardle, Serge Mostowy

**Author notes:** Correspondence (M.C.G.), (S.M.).

## Abstract

Trained immunity is a long-term memory of innate immune cells, generating an improved response upon re-infection. *Shigella* is an important human pathogen and inflammatory paradigm for which there is no effective vaccine. Using zebrafish larvae we demonstrate that after *Shigella* priming neutrophils are more efficient at bacterial clearance. We observe that *Shigella*-induced protection is non-specific and long-lasting, and is unlike training by BCG and β-glucan. Analysis of histone ChIP-seq on primed neutrophils revealed that *Shigella* training deposits the active H3K4me3 mark on promoter regions of 1612 genes, significantly changing the epigenetic landscape of neutrophils towards enhanced microbial recognition and mitochondrial ROS production. Finally, we demonstrate that mitochondrial ROS plays a key role in enhanced antimicrobial activity of trained neutrophils. It is envisioned that signals and mechanisms we discover here can be used in other vertebrates, including humans, to suggest new therapeutic strategies involving neutrophils to control bacterial infection.

## INTRODUCTION

Trained immunity is an immunological memory characterized by remodelling of the epigenetic landscape and metabolism of myeloid cells, conferring enhanced responses upon a new infection challenge ^1^. The most commonly used training stimuli are *Mycobacterium bovis* Bacille Calmette-Guérin (BCG) and the fungal wall component β-glucan ^1–5^, however reports have described a large breadth of triggers that include metabolites, inflammatory cytokines and danger signals ^2,3,6–9^. Although work *in vitro, ex vivo*, in mice and in humans has mostly been centred around monocytes/macrophages ^1,10,11^, growing evidence shows that mechanisms of trained immunity can be observed in other immune and non-immune cells (such as endothelial and epithelial cells) ^12–16^. Recent reports have started to investigate training mechanisms by BCG and β-glucan in neutrophils ^17,18^, however the impact of trained neutrophils at a whole animal level is mostly unknown.

The zebrafish embryo has been widely used as a model for developmental biology, cell biology and infectious diseases ^19–22^. Zebrafish are genetically tractable, share >80% of human genes associated with diseases, and strictly depend upon innate immune responses until 4 weeks post-fertilization when the adaptive immune system starts to develop ^23–25^. Seminal studies have used zebrafish as a model to study hematopoietic stem and progenitor cells (HSPCs) development and haematopoiesis ^26–30^, and the repertoire of innate immune cells in zebrafish is now well characterized ^31,32^. Considering this, the zebrafish model presents itself as an attractive model to study trained immunity. The first evidence that training mechanisms may also be conserved in fish comes from decades of use of β-glucans in aquaculture, where their use improved the resistance of teleost fish to infections ^33,34^. Further work with carp macrophages ^35^, vaccine development ^36–38^ and pathogenic infections ^29,39,40^ support the use of fish as a model to dissect fundamental determinants of trained immunity.

*Shigella* is an important human pathogen, among the top 12 priority pathogens requiring urgent action by the World Health Organization ^41^, and for which there are no effective vaccines ^42^. *Shigella* infection is well known to induce inflammation, by its lipopolysaccharide (LPS) and type III secretion system (T3SS) ^43,44^. We previously demonstrated that hallmarks of *Shigella* infection, including inflammation and macrophage cell death, can be observed using a zebrafish infection model ^45–48^. Work has also shown that a non-lethal *Shigella* infection triggers immune responses that lead to emergency granulopoiesis, and protects zebrafish larvae from secondary infection ^40,49^.

Here we introduce the zebrafish larvae infection model as a powerful system to study innate immune training. We show that *Shigella*-priming of zebrafish larvae generates protective neutrophils, and the training mechanism differs from that of BCG- or β-glucan-priming. Genome-wide sequencing of epigenetic modifications in trimethylation at histone 3 lysine 4 (H3K4me3) in *Shigella*-primed neutrophils and downstream functional characterisation shows that their epigenetic landscape is modified to increase mitochondrial reactive oxygen species (mtROS) production to better fight secondary infectious challenges.

## RESULTS

### Zebrafish priming with a non-lethal dose of *Shigella* induces generation of protective neutrophils

We previously demonstrated that a non-lethal dose of *Shigella flexneri* injected in the hindbrain ventricle (HBV) of zebrafish larvae at 2 days post-fertilisation (dpf) is rapidly cleared and induces hematopoietic stem and progenitor cell (HSPC) proliferation and differentiation as well as emergency granulopoiesis in the aorta-gonad-mesonephros (AGM) region ^40^. To better understand the immune responses during a non-lethal *Shigella* infection, we quantified total number of neutrophils and macrophages in larvae injected with ~2×10^3^ CFU (Fig 1A). Neutrophil numbers significantly decrease at 24 hours post 1^st^ infection (hp1i), and by 48 hp1i the pool of neutrophils is replenished in excess, consistent with observed granulopoiesis in the AGM (Fig 1B-C). Although the macrophage population is also significantly reduced by 24 hp1i, it does not recover by 48 hp1i (Fig 1D), suggesting that macrophage hematopoiesis is impaired to favor granulopoiesis, consistent with previous reports ^2,8,50,51^.

**Figure 1.**
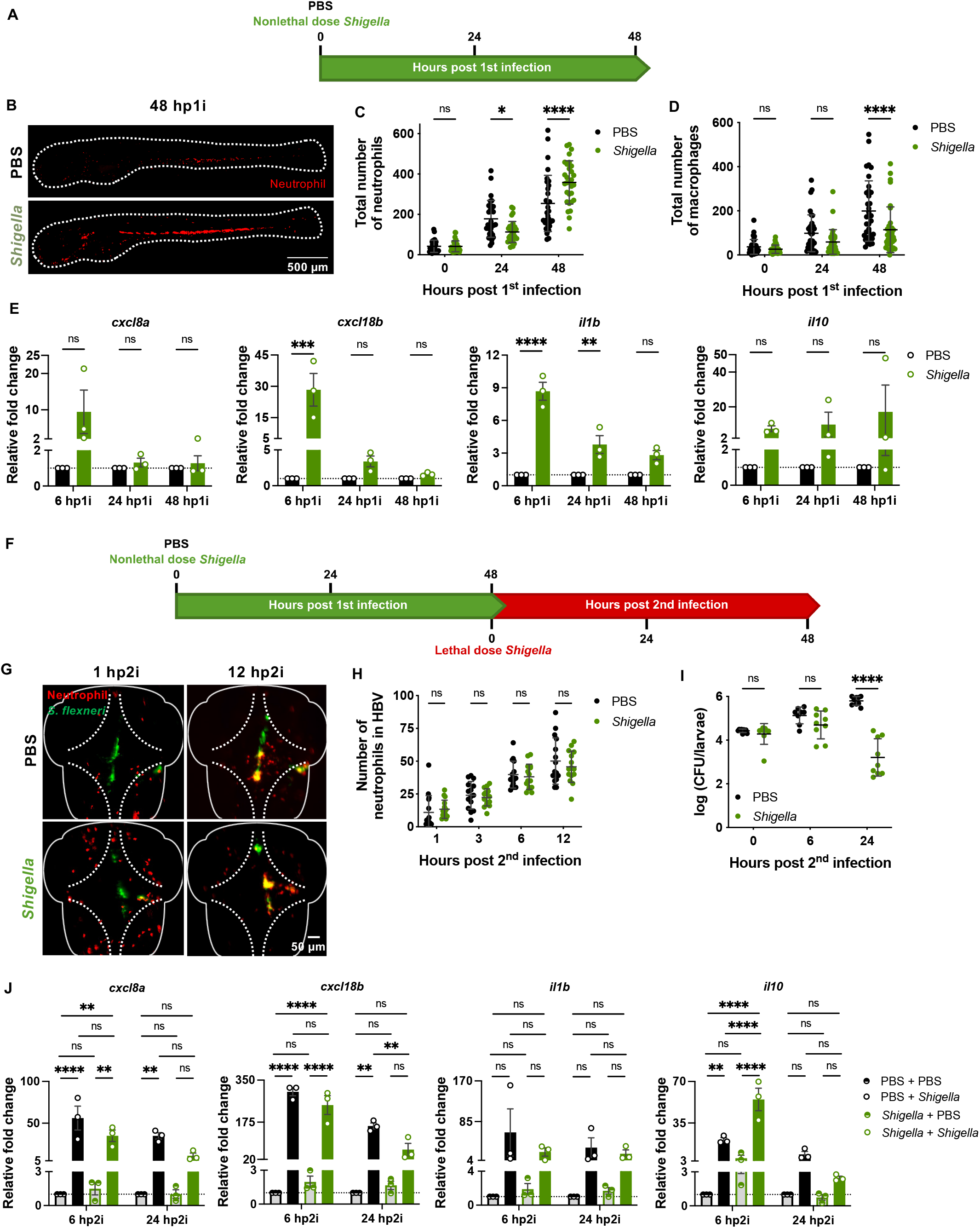
Zebrafish exposure to non-lethal *Shigella* dose induces generation of more resilient neutrophils. **A** Experimental outline of 2 dpf zebrafish larvae injected in the HBV with PBS or a non-lethal dose of *Shigella*. **B** Representative images of Tg(*lyz*::dsRed) larvae with red neutrophils at 48 hp1i with a non-lethal dose of *S. flexneri* M90T. Scale bar: 500 μm. **C** Quantification of neutrophils (Tg(*lyz*::dsRed)) at 0, 24 and 48 hp1i injected with PBS (naïve-black full circles) and a non-lethal dose of *S. flexneri* M90T (green full circles, 2×10^3^ ± 1.4×10^3^ CFUs). Data pooled from 3 experiments with n=10 larvae per time point per condition per experiment (mean ± SD, *p < 0.05, ***p < 0.001, 2-way ANOVA with Sidak’s multiple comparisons test). **D** Quantification of macrophages (Tg(*mpeg*::mCherry)) at 0, 24 and 48 hp1i injected with PBS (naïve-black full circles) and a non-lethal dose of *S. flexneri* M90T (green full circles, 1.8×10^3^ ± 8.6×10^2^ CFUs). Data pooled from 3 experiments with n>11 larvae per time point per condition per experiment (mean ± SD, ****p < 0.0001, 2-way ANOVA with Sidak’s multiple comparisons test). **E** Fold change in the expression of *cxcl8a*, *cxcl18b*, *il1b* and *il10* in larvae injected with non-lethal dose of *S. flexneri* M90T (green full bars, 1.3×10^3^ ± 5.5×10^2^ CFUs) as compared to PBS control (naïve-black full bars). Data pooled from 3 experiments with n>5 larvae per time point per condition per experiment (mean ± SEM, **p < 0.01, ***p < 0.001, ****p < 0.0001, 2-way ANOVA with Sidak’s multiple comparisons test). **F** Experimental outline of 2 dpf zebrafish larvae injected in the HBV with PBS or a non-lethal dose of *Shigella* followed by a lethal dose injection of *S. flexneri* M90T at 4dpf in the HBV. **G** Representative images of naïve and *Shigella*-primed larvae (Tg(*lyz*::dsRed)) with red neutrophils infected with a lethal dose of *S. flexneri* M90T at 1 and 12 hp2i. Scale bar: 50 μm. **H** Quantification of recruited neutrophils (Tg(*lyz*::dsRed)) to the HBV in naïve (black full circles) and *Shigella*-primed (green full circles) larvae at 1, 3, 6 and 12 h following injection of a lethal dose of *S. flexneri* M90T (PBS - 2.6×10^4^ ± 9×10^3^ CFUs, *Shigella* - 2.7×10^4^ ± 1×10^4^ CFUs). Data pooled from 3 experiments with n=5 larvae per time point per condition per experiment (mean ± SD, 2-way ANOVA with Sidak’s multiple comparisons test). **I** Log_10_-transformed CFU counts of naïve and *Shigella*-primed larvae injected with a lethal dose of *S. flexneri* M90T (PBS - 2.7×10^4^ ± 4.5×10^3^ CFUs, *Shigella* - 2.5×10^4^ ± 1×10^4^ CFUs). Data pooled from 3 independent experiments using n=3 larvae per condition per experiment (mean ± SD, ****p < 0.0001, 2-way ANOVA with Sidak’s multiple comparisons test). **J** Fold change in the expression of *cxcl8a, cxcl18b, il1b* and *il10* following lethal dose injection of *S. flexneri* M90T in naïve (black full bars, 2.5×10^4^ ± 1×10^4^ CFUs) and *Shigella*-primed (green full bars, 2.7×10^4^ ± 4.5×10^3^ CFUs) as compared to PBS injected controls (grey bars). Data pooled from 3 experiments with n>5 larvae per time point per condition per experiment (mean ± SEM, **p < 0.01, ****p < 0.0001, 2-way ANOVA with Sidak’s multiple comparisons test).

Analysis of cytokine expression showed a significant increase in pro-inflammatory cytokines *cxcl8a, cxcl18b, il1b, il6* and *tnfa* between 6 and 24 hp1i (Fig 1E and Fig S1A), which correlates with neutrophil recruitment and bacterial clearance (Fig S1B). At 48 hp1i, the pro-inflammatory response induced by *Shigella* priming returns to basal levels, but expression of the anti-inflammatory cytokine *il10* remains elevated. These results are consistent with the requirement of inflammation to induce proliferation of HSPCs ^28,29,40^, and show that primed larvae (*i.e*. those injected with a non-lethal dose of *Shigella*) are at immunological homeostasis prior to reinfection.

Upon a 2^nd^ lethal *Shigella* challenge (Fig 1F), primed larvae show significantly higher survival rates by 48 hours post 2^nd^ infection (hp2i) as compared to naïve larvae (*i.e*. those injected with PBS). Quantification of neutrophil recruitment to the HBV when infected with a lethal dose (>2×10^4^ CFU), showed that recruitment in *Shigella*-primed larvae is not significantly different from naïve larvae (Fig 1G-H). At 6 hp2i, bacterial burden in *Shigella*-primed larvae starts to reduce, as compared to naïve larvae (Fig 1I), suggesting that primed neutrophils are better at clearing bacteria. Similarities in neutrophil recruitment in both conditions during infection are consistent with expression of pro-inflammatory cytokines *cxcl8a* and *cxcl18b*, which are expressed to the same extent in naïve and primed larvae at 6 hp2i (Fig 1J). In primed infected larvae, the anti-inflammatory cytokine *il10* expression is significantly different compared to that in naïve larvae (both uninfected and infected), suggesting that primed larvae strongly induce anti-inflammatory responses. At 24 hp2i, all analysed cytokines are less expressed in primed infected larvae compared to naïve infected larvae (Fig 1J and Fig S1B). Together, these observations indicate that *Shigella*-primed larvae are protected against a secondary infectious challenge because of tighter control of immune responses and their neutrophils have increased killing capacity, which are important characteristics of innate immune training.

### *Shigella* innate immune training is non-specific and long lasting

Work on trained innate immunity using BCG has demonstrated that induced protection is not specific to *Mycobacterium tuberculosis* (Mtb) and helps in controlling fungal infection by *Candida albicans* ^52^. In zebrafish, β-glucan and *Salmonella enterica* serovar Typhimurium have been shown to protect against Gram-negative and Gram-positive pathogens ^39^. Here we tested our *Shigella*-primed larvae against lethal infections of *Pseudomonas aeruginosa* (a Gram-negative pathogen, 2×10^3^ CFU), and *Staphylococcus aureus* (a Gram-positive pathogen, 2×10^4^ CFU) (Fig 2A). In both cases, the survival of *Shigella*-primed larvae is significantly higher as compared to the survival of naïve larvae (Fig 2B-C), and bacterial burden is controlled from 6 hp2i (Fig S2A-B). These results demonstrate that *Shigella*-induced protection of zebrafish is non-specific.

**Figure 2.**
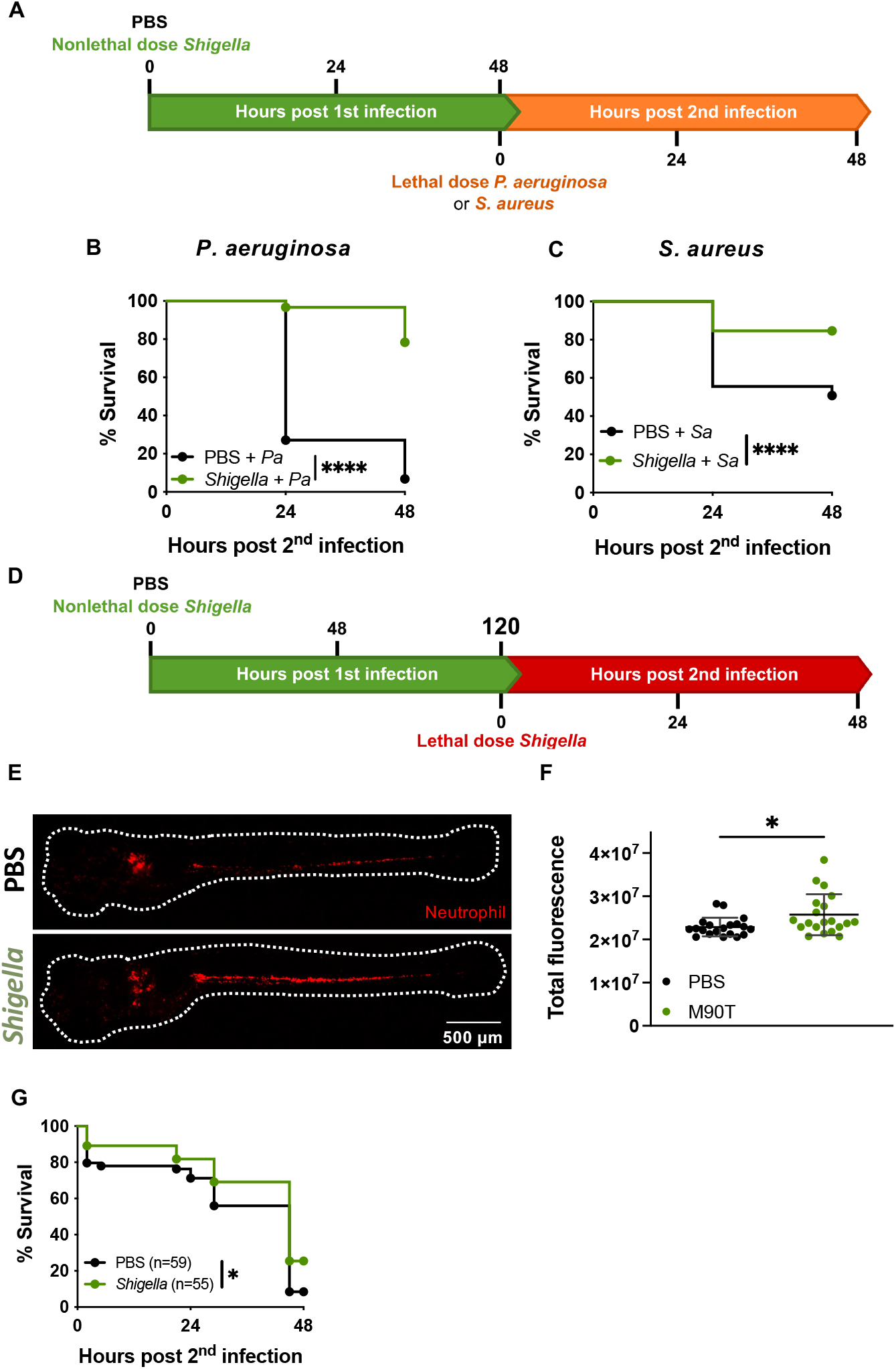
*Shigella* induced protection is non-specific and long lasting. **A** Experimental outline of 2 dpf zebrafish larvae injected in the HBV with PBS or a non-lethal dose of *Shigella* followed by a lethal dose injection of *P*. aeruginosa or *S. aureus* at 4 dpf in the HBV. **B** Survival curves from naïve (black full circles) and *Shigella*-primed (green full circles, 2.2×10^3^ ± 3.8×10^2^ CFUs) larvae infected with a lethal dose of *P. aeruginosa* (PBS – 2.6×10^3^ ± 7.4×10^2^ CFUs, *Shigella* – 2.3×10^3^ ± 5.5×10^2^ CFUs). Data pooled from 3 independent experiments using n=20 larvae per condition per experiment (****p < 0.0001, Log-rank (Mantel-Cox) test). **C** Survival curves from naïve (black full circles) and *Shigella*-primed (green full circles, 1.8×10^3^ ± 7×10^2^ CFUs) larvae infected with a lethal dose of *S. aureus* (PBS – 1.7×10^4^ ± 4.5×10^3^ CFUs, *Shigella* – 2.6×10^4^ ± 1×10^4^ CFUs). Data pooled from 3 independent experiments using n> 20 larvae per condition per experiment (****p < 0.0001, Log-rank (Mantel-Cox) test). **D** Experimental outline of 2 dpf zebrafish larvae injected in the HBV with PBS or a non-lethal dose of *Shigella* followed by a lethal dose injection of *S. flexneri* M90T at 7 dpf (or 5 dpi) in the HBV. **E** Representative images of naïve and *Shigella*-primed Tg(*lyz*::dsRed) larvae with red neutrophils at 5 dp1i. **F** Quantification of total fluorescence in Tg(*lyz*::dsRed) naïve (black full circles) and *Shigella*-primed (green full circles, 1.7×10^3^ ± 5.7×10^2^ CFUs) larvae at 5dp1i. Data pooled from 2 experiments with n=10 larvae per condition per experiment (mean ± SD, *p < 0.05, unpaired Student’s *t*-test). **G** Survival curves from naïve (black full circles) and *Shigella*-primed (green full circles) larvae infected with a lethal dose of *S. flexneri* M90T at 5 dp1i. Data pooled from 3 independent experiments using n>16 larvae per condition per experiment (*p < 0.05, Logrank (Mantel-Cox) test). **H** Log_10_-transformed CFU counts of naïve and *Shigella*-primed (in green section: 1.7×10^3^ ± 5.7×10^2^ CFUs) larvae injected with a lethal dose of *S. flexneri* M90T (in red section: PBS - 2.7×10^4^ ± 4.5×10^3^ CFUs, *Shigella* - 2.5×10^4^ ± 1×10^4^ CFUs). Data pooled from 3 independent experiments using n>2 larvae per condition per experiment (mean ± SD, 2-way ANOVA with Sidak’s multiple comparisons test).

To test the longevity of protection induced by *Shigella* priming, we increased the resting interval between priming and reinfection from 2 to 5 days (Fig 2D). At this timepoint (7 dpf), we observed that neutrophil numbers remain higher in primed larvae as compared to naïve larvae (Fig 2E-F), but the differences are smaller than at 4 dpf (Fig 1B-C). Upon reinfection with a lethal dose of *Shigella* (2.5×10^4^ CFU), survival of primed larvae is significantly higher compared to naïve larvae (Fig 2G). However, differences in bacterial clearance are less detectable at 24 hp2i between groups (Fig S2C).

### BCG and β-glucan induce protection in zebrafish larvae

BCG and β-glucan have been widely studied as inducers of trained immunity ^52,53^. To better understand the mechanisms by which *Shigella* can induce trained immunity, we developed zebrafish reinfection models with BCG and β-glucan for comparison. In the BCG reinfection model (Fig 3A), a low dose of BCG (50 CFU) was injected in 2 dpf larvae that were then incubated at 33 ^o^C for 48 h to stimulate BCG replication ^54^. Although BCG were not completely cleared from larvae (Fig 3C), reinfection with a lethal dose of *Shigella* was done at 48 hp1i and survival was followed for another 48 h at 28.5 ^o^C. In this case, BCG-primed larvae had significantly greater survival than naïve larvae (Fig 3B), even though they presented a similar burden of *Shigella* at 24 hp2i (Fig 3C). The difference between BCG and *Shigella* training may be associated with pro-inflammatory responses that are induced during priming. Whereas *Shigella* strongly induces pro-inflammatory responses by 6 hp1i (Fig 1E), BCG infection does not (Fig 3D). Surprisingly, the expression of neutrophil chemokines *cxcl8a* and *cxcl18b* were repressed at 6 hp1i, which may explain the lack of neutrophil recruitment to the HBV by 24 hp1i (Fig S3A). Macrophages, in contrast, were recruited to the injection site and remained in the HBV until 24 hp1i (Fig S3B). Overall, no differences in neutrophil and macrophage numbers were observed during BCG infection (Fig 3E-G).

**Figure 3.**
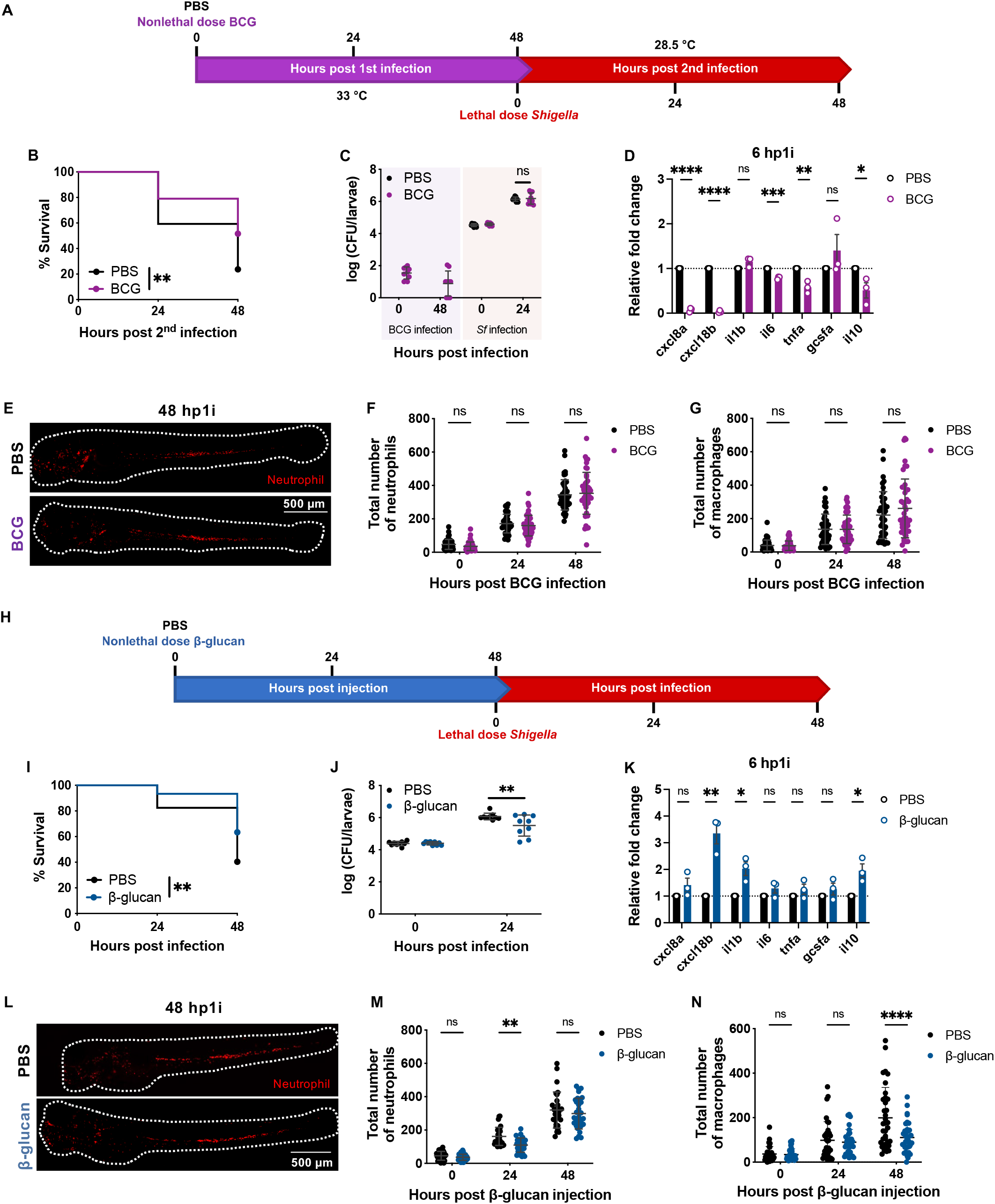
BCG and β-glucan induce protection in zebrafish embryos. **A** Experimental outline of 2 dpf zebrafish larvae injected in the HBV with PBS or a non-lethal dose of BCG followed by a lethal dose injection of *S. flexneri* M90T at 4 dpf in the HBV. During priming (48 hp1i) larvae were kept at 33°C and during reinfection they were kept at 28.5°C. **B** Survival curves from naïve (black full circles) and BCG-primed (purple full circles) larvae infected with a lethal dose of *S. flexneri* M90T. Data pooled from 3 independent experiments using n>17 larvae per condition per experiment (**p < 0.01, Log-rank (Mantel-Cox) test). **C** Log_10_-transformed CFU counts of naïve and BCG-primed (in purple section: 4.7×10^1^ ± 3.2×10^1^ CFUs) larvae injected with a lethal dose of *S. flexneri* M90T (in red section: PBS – 3.5×10^4^ ± 6.8×10^3^ CFUs, BCG – 3.9×10^4^ ± 7.5×10^3^ CFUs). Data pooled from 3 independent experiments using n=3 larvae per condition per experiment (mean ± SD, 2-way ANOVA with Sidak’s multiple comparisons test). **D** Fold change in the expression of *cxcl8a, cxcl18b, il1b, il6, tnfa, gcsfa* and *il10* following non-lethal dose injection of BCG (purple bars, 4.7×10^1^ ± 3.2×10^1^ CFUs) as compared to PBS injected controls (grey bars). Data pooled from 3 experiments with n=10 larvae per time point per condition per experiment (mean ± SEM, *p < 0.05, **p < 0.01, ***p < 0.001, ****p < 0.0001, unpaired Student’s *t*-test). **E** Representative images of naïve and BCG-primed Tg(*lyz*::dsRed) larvae with red neutrophils at 48 hp1i. Scale bar: 500 μm. **F** Quantification of neutrophils (Tg(*lyz*::dsRed)) at 0, 24 and 48 hp1i injected with PBS (naïve-black full circles) and a non-lethal dose of BCG (purple full circles, 5.6×10^1^ ± 4.3×10^1^ CFUs). Data pooled from 3 experiments with n>13 larvae per time point per condition per experiment (mean ± SD, 2-way ANOVA with Sidak’s multiple comparisons test). **G** Quantification of macrophages (Tg(*mpeg*::mCherry)) at 0, 24 and 48 hp1i injected with PBS (naïve-black full circles) and a non-lethal dose of BCG (purple full circles, 6.3×10^1^ ± 3.7×10^1^ CFUs). Data pooled from 3 experiments with n>12 larvae per time point per condition per experiment (mean ± SD, 2-way ANOVA with Sidak’s multiple comparisons test). **H** Experimental outline of 2 dpf zebrafish larvae injected in the HBV with PBS or non-lethal dose of β-glucan followed by a lethal dose injection of *S. flexneri* M90T at 4 dpf in the HBV. **I** Survival curves from naïve (black full circles) and β-glucan-primed (blue full circles, 20 ng) larvae infected with a lethal dose of *S. flexneri* M90T. Data pooled from 3 independent experiments using n>18 larvae per condition per experiment (**p < 0.01, Log-rank (Mantel-Cox) test). **J** Log_10_-transformed CFU counts of naïve and β-glucan-primed larvae injected with a lethal dose of *S. flexneri* M90T (PBS – 2.5×10^4^ ± 7.3×10^3^ CFUs, β-glucan – 2.6×10^4^ ± 5.9×10^3^ CFUs). Data pooled from 3 independent experiments using n=3 larvae per condition per experiment (mean ± SD, **p < 0.01, 2-way ANOVA with Sidak’s multiple comparisons test). **K** Fold change in the expression of *cxcl8a, cxcl18b, il1b, il6, tnfa, gcsfa* and *il10* following non-lethal dose injection of β-glucan (blue bars, 20 ng) as compared to PBS injected controls (black bars). Data pooled from 3 experiments with n=10 larvae per time point per condition per experiment (mean ± SEM, *p < 0.05, **p < 0.01, unpaired Student’s *t*-test). **L** Representative images of naïve and β-glucan-primed Tg(*lyz*::dsRed) larvae with red neutrophils at 48 hp1i. Scale bar: 500 μm. **M** Quantification of neutrophils (Tg(*lyz*::dsRed)) at 0, 24 and 48 hp1i injected with PBS (naïve-black full circles) and a non-lethal dose of β-glucan (blue full circles, 20 ng). Data pooled from 3 experiments with n>8 larvae per time point per condition per experiment (mean ± SD, **p < 0.01, 2-way ANOVA with Sidak’s multiple comparisons test). **N** Quantification of macrophages (Tg(*mpeg*::mCherry)) at 0, 24 and 48 hp1i injected with PBS (naïve-black full circles) and a non-lethal dose of β-glucan (blue full circles, 20 ng). Data pooled from 3 experiments with n>11 larvae per time point per condition per experiment (mean ± SD, ****p < 0.0001, 2-way ANOVA with Sidak’s multiple comparisons test).

To establish a β-glucan (local) reinfection model (Fig 3H), 20 ng of β-glucan were injected in the HBV of larvae and infection with *Shigella* occurred 48 h after. Larvae primed with β-glucan had increased survival (Fig 3I) and reduced bacterial burden by 24 hp2i (Fig 3J). Analysis of cytokine expression at 6 hp1i demonstrated that *cxcl18b, il1b* and *il10* are significantly more expressed in β-glucan primed larvae compared to naïve larvae (Fig 3K), suggesting that β-glucan is recognized by the immune system. Similar to *Shigella* priming, β-glucan induced changes in the neutrophil and macrophage populations. At 24 hp1i the neutrophil numbers were significantly reduced, and by 48 hp1i the population was restored to similar numbers as in naïve larvae (Fig 3L-M). The granulopoiesis observed was associated to expression of *gcsfa* expression at 24 hp1i (Fig S3F), but there was no significant increase in the number of neutrophils in the AGM as compared to naïve larvae (Fig S3G). Consequently, and similar to *Shigella* priming, macrophage differentiation was blunted by 48 hp1i (Fig 3N).

Considering the differences across zebrafish models of trained innate immunity (Table 1), we conclude that the priming mechanism induced by *Shigella* is different from that of BCG and β-glucan.

**Table 1.** Comparison of training phenotypes between *Shigella*-, BCG- and β-glucan-primed zebrafish larvae.

| | | *Shigella* | BCG | β-glucan |
| --- | --- | --- | --- | --- |
| **Training** | Bacterial clearance | YES | NO | -- |
|  | Neutropenia (24 hp1i) | YES | NO | YES |
|  | Emergency granulopoiesis (48 hp1i) | YES | NO | NO |
|  | Changes in macrophage population | YES | NO | YES |
|  | Induction pro-inflammatory responses | YES | NO | YES |
| **Infection challenge** | Survival protection | YES | YES | YES |
|  | Infection control | YES | NO | YES |
|  | Non-specific | YES | YES | YES |

### Contribution of host factors, LPS and *Shigella* effectors to training mechanisms

*Gcsf* signalling is crucial to induce emergency granulopoiesis in *Shigella*-primed larvae ^40^, however it is not the only requirement to induce training mechanisms. To better understand triggers for *Shigella*-induced training in zebrafish larvae, we tested different factors important in *Shigella* infection of host cells (Fig 3).

As previously described ^28^, inflammasome activation plays an important role in HSPCs production and *Shigella* has been shown to be a potent activator of inflammasome pathways ^43,44^. Therefore, we tested if NLRP3 inflammasome activation by nigericin and glucose in HSPCs is sufficient to induce training. Zebrafish larvae were bathed in either 0.1 μM nigericin or 1% glucose for at least 24 h from 2 dpf, and at 4 dpf larvae were injected with a lethal dose of *Shigella* (Fig 4A). The survival assay showed that independently of exposure time to either nigericin or glucose, no differences in survival (Fig 4B-C) or bacterial burden (Fig S4A-B) are observed. These results suggest that inflammasome and caspase-1 activation is not sufficient to induce training.

**Figure 4.**
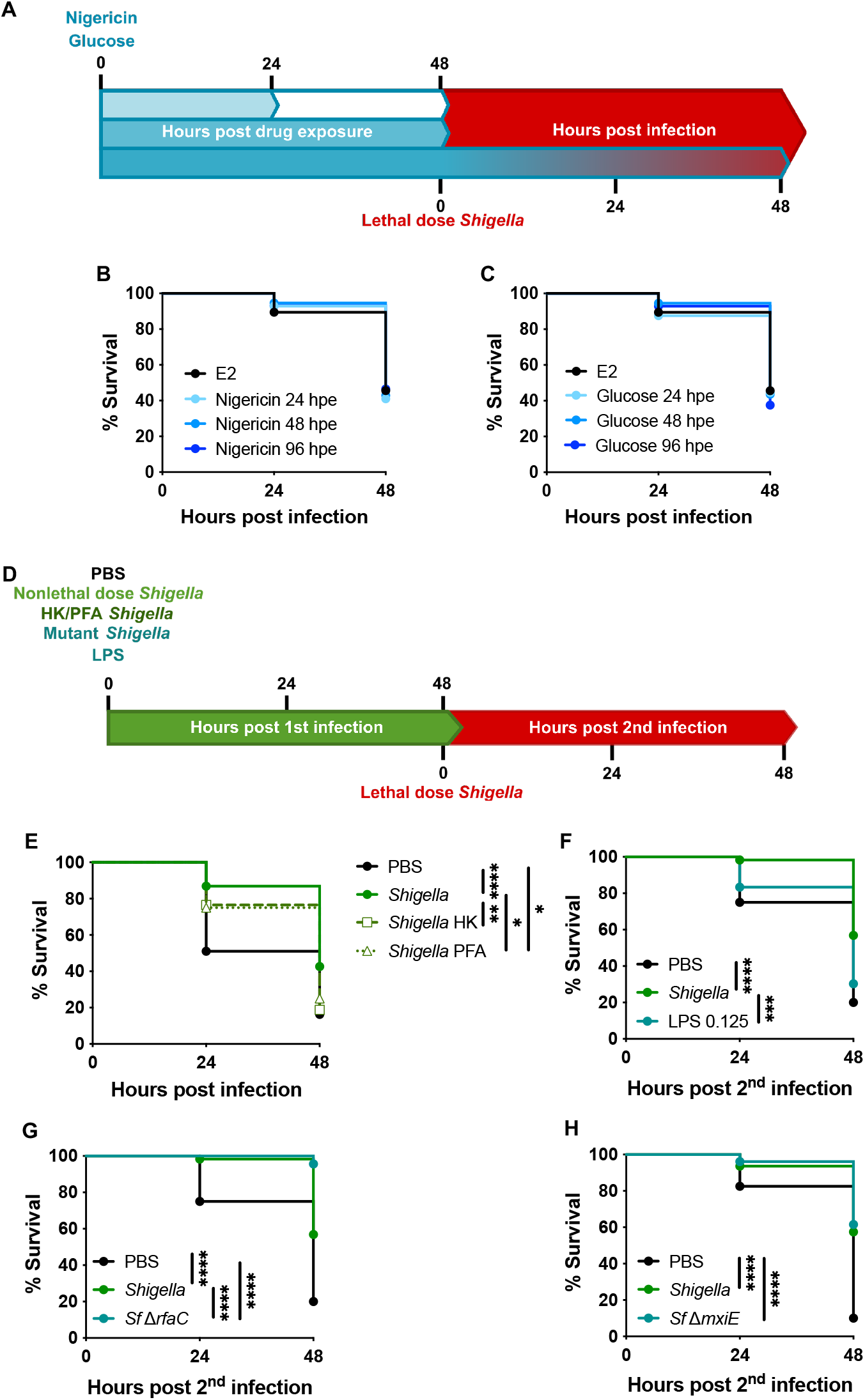
*Shigella* stimulates inflammatory pathways to induce training. **A** Experimental outline of exposure of 2 dpf zebrafish larvae to nigericin and glucose for 24, 48 or 96h followed by a lethal dose injection of *S. flexneri* M90T at 4 dpf in the HBV. **B** Survival curves from E2 (embryo media control, black full circles) and nigericin (blue full circles, 0.1 μM) treated larvae infected with a lethal dose of *S. flexneri* M90T (PBS – 2.7×10^4^ ± 8.5×10^3^ CFUs, nigericin – 2.6×10^4^ ± 4.1×10^3^ CFUs). Data pooled from 3 independent experiments using n>14 larvae per condition per experiment (Log-rank (Mantel-Cox) test). **C** Survival curves from E2 (embryo media control, black full circles) and glucose (blue full circles, 1%) treated larvae infected with a lethal dose of *S. flexneri* M90T (PBS – 2.7×10^4^ ± 8.5×10^3^ CFUs, glucose – 2.7×10^4^ ± 1×10^4^ CFUs). Data pooled from 3 independent experiments using n>14 larvae per condition per experiment (Log-rank (Mantel-Cox) test). **D** Experimental outline of 2 dpf zebrafish larvae injected in the HBV with PBS, non-lethal dose of live or killed *Shigella, Shigella* mutants or LPS followed by a lethal dose injection of *S. flexneri* M90T at 4 dpf in the HBV. **E** Survival curves from naïve (black full circles), live *Shigella*-primed (green full circles, 1.3×10^3^ ± 6.6×10^2^ CFUs), heat-killed (HK) *Shigella*-primed (dark green/white squares) and PFA-killed *Shigella*-primed (dark green/white triangles) larvae infected with a lethal dose of *S. flexneri* M90T (PBS – 2.7×10^4^ ± 6.1×10^3^ CFUs, *Shigella* – 2.7×10^4^ ± 1×10^4^ CFUs, *Shigella* HK – 3.2×10^4^ ± 3.3×10^4^ CFUs, *Shigella* PFA – 3.2×10^4^ ± 1.7×10^4^ CFUs). Data pooled from 3 independent experiments using n>11 larvae per condition per experiment. (*p < 0.05, **p < 0.01, ****p < 0.0001, Log-rank (Mantel-Cox) test). **F** Survival curves from naïve (black full circles), *Shigella*-primed (green full circles, 1.6×10^3^ ± 6.5×10^2^ CFUs) and LPS-primed (moss green full circles, 0.125 ng) larvae infected with a lethal dose of *S. flexneri* M90T (PBS – 1.9×10^4^ ± 6.3×10^3^ CFUs, *Shigella* – 2.8×10^4^ ± 4.7×10^3^ CFUs, LPS – 1.9×10^4^ ± 9.1×10^3^ CFUs). Data pooled from 3 independent experiments using n>18 larvae per condition per experiment. (***p < 0.001, ****p < 0.0001, Log-rank (Mantel-Cox) test). **G** Survival curves from naïve (black full circles), *Shigella*-primed (green full circles, 1.6×10^3^ ± 6.5×10^2^ CFUs) and Shigella Δ*rfaC*-primed (moss green full circles, 8.5×10^2^ ± 7.5×10^2^ CFUs) larvae infected with a lethal dose of *S. flexneri* M90T (PBS – 1.9×10^4^ ± 6.3×10^3^ CFUs, *Shigella* – 2.8×10^4^ ± 4.7×10^3^ CFUs, Δ*rfaC* – 2.4×10^4^ ± 8.4×10^3^ CFUs). Data pooled from 3 independent experiments using n>10 larvae per condition per experiment (****p < 0.0001, Log-rank (Mantel-Cox) test). **H** Survival curves from naïve (black full circles), *Shigella*-primed (green full circles, 7.9×10^2^ ± 5.5×10^2^ CFUs) and Shigella Δ*mxiE*-primed (moss green full circles, 8.8×10^2^ ± 5.8×10^2^ CFUs) larvae infected with a lethal dose of *S. flexneri* M90T (PBS – 1.9×10^4^ ± 7.5×10^3^ CFUs, *Shigella* – 2.5×10^4^ ± 4.6×10^3^ CFUs, Δ*mxiE* – 2.2×10^4^ ± 7.6×10^3^ CFUs). Data pooled from 3 independent experiments using n>10 larvae per condition per experiment. (****p < 0.0001, Log-rank (Mantel-Cox) test).

Recently, studies with BCG and *Salmonella* have shown that inactivated or heat-killed bacteria can induce trained immunity mechanisms, however the protection observed upon second challenge is weaker ^39,54,55^. In the case of *Shigella* (Fig 4D), both heat-killed and PFA-killed *Shigella* failed to protect larvae survival upon a secondary lethal dose of *Shigella* (Fig 4E and Fig S4C), strongly suggesting that *Shigella* needs to be alive to induce training.

To test the role of the lipopolysaccharide (LPS) in training, LPS was used to prime larvae followed by secondary challenge with a lethal dose of *Shigella* (Fig 4F and Fig S4D-F). Larvae survival upon a lethal *Shigella* injection was not significantly affected by priming with LPS, highlighting that in our model LPS alone does not induce protection mechanisms. However, complete exposure of lipid A in a *Shigella* Δ*rfaC* mutant (which lacks the O-antigen, outer and inner core of LPS, ^56^) induced 100% protection upon reinfection with *Shigella* (Fig 4G) and highly efficient bacterial clearance (Fig S4G). Virulence assays in zebrafish larvae showed that 1×10^4^ CFU of *Shigella rfaC* mutant can cause 100% lethality in 48 hpi, whereas *Shigella* wild-type (WT) only kills 50% of the larvae (Fig S4H). The lethality was found to be caused by heighted pro-inflammatory responses, and not due to bacterial burden (Fig S4I-J). These results support work showing that stronger pro-inflammatory responses induced during priming can result in better innate immune training ^54^.

Previous work has shown that the T3SS plays a role in training, where a complete and functional T3SS provides more protection than an absent T3SS and its effectors ^40^. Here we tested the effect of T3SS effectors regulated by the master regulator MxiE ^57^. Larvae survival upon reinfection showed that *mxiE* regulated effectors do not significantly impact how *Shigella* induces training (Fig 4H and Fig S4K). We also tested a *Shigella ospF* mutant, lacking the bacterial immunomodulin OspF ^58–60^, where similar results to *mxiE* mutant were obtained (Fig S4L-M). These results are consistent with similar levels of virulence observed during infection of zebrafish larvae with *Shigella* Δ*mxiE* and Δ*ospF* mutants and *Shigella* WT (Fig S4N-Q).

Together these data suggest that live *Shigella*, with a functional T3SS, is required to induce inflammatory responses that trigger training mechanisms.

### Exposure to *Shigella* induces epigenetic reprogramming of neutrophils

Epigenetic reprogramming is a defining characteristic underlying the innate immune training of myeloid cells and the enrichment of trimethylation of lysine 4 at histone 3 (H3K4me3) on gene promoter regions is typically associated to active/poised gene transcription ^1^. Considering this, we investigated if *Shigella*-primed neutrophils exhibit different H3K4me3 patterns that could explain their ability to better respond to reinfection. To test this, neutrophils were FACS sorted from naïve and primed (*Shigella* WT and *Shigella* ΔT3SS (i.e. *mxiD* mutant)) larvae at 48 hp1i (Fig S5A) and chromatin immunoprecipitation sequencing (ChIP-seq) was performed on the H3K4me3 mark (Fig 5A). Genes were then annotated if they had a H3K4me3 mark in their promoter region (defined as ±3kb from the transcription start site; Tables S1-S3). Naïve neutrophils displayed H3K4me3 peaks at the promoter region of 1049 genes, which are associated with normal cell functioning (Fig S5B-C), including surveying the quality of mRNA to increase the fidelity of RNA processing. Strikingly, neutrophil samples primed with *Shigella* WT or ΔT3SS showed an increase in the number of genes marked by H3K4me3 (2242 and 2707 respectively) as compared to naïve neutrophils, suggesting epigenetic remodelling of primed neutrophils (Fig 5B and C, Fig S5D-E).

**Figure 5.**
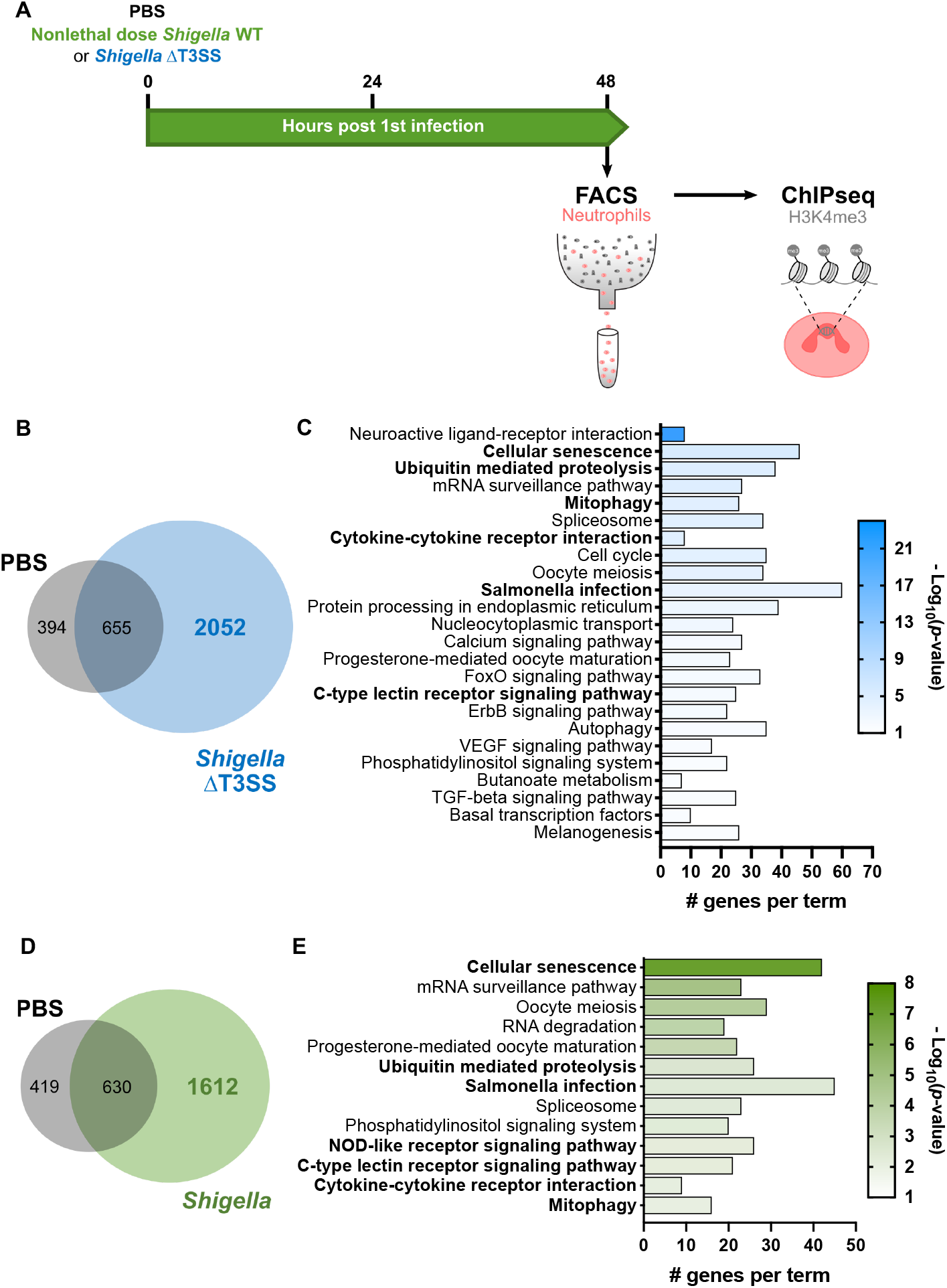
Exposure to *Shigella* induces epigenetic reprogramming of neutrophils. **A** Experimental outline of FACS-sorting neutrophils from naïve and *Shigella*-primed larvae for ChIP-seq on H3k4me3 modification. **B** Venn diagram showing number of common and unique genes marked by H3K4me3 peaks in their promoter regions (±3kb from TSS) of PBS and *Shigella* ΔT3SS primed larvae. **C** Enriched KEGG-pathways (*p*-value < 0.01) associated with the 2052 unique *Shigella* ΔT3SS priming genes marked by a H3K4me3 peak in their promoter regions (±3kb from TSS). **D** Venn diagram showing number of common and unique genes marked by H3K4me3 peaks in their promoter regions (±3kb from TSS) of PBS and *Shigella* WT primed larvae. **E** Enriched KEGG-pathways (*p*-value < 0.01) associated with the 1612 unique *Shigella* WT priming genes marked by a H3K4me3 peak in their promoter regions (±3kb from TSS).

Upon *Shigella* ΔT3SS priming, pathway analysis of marked gene promoter regions indicates that some modulated responses are common to *Shigella* WT-priming, however many other different signalling pathways are affected (Fig 5C and Table S2), highlighting that priming with *Shigella* WT leads to a more precise and efficient immune response. *Shigella* WT induced H3K4me3 deposition at promoter regions was observed for genes involved in cellular senescence and in infection signalling pathways, specifically associated to infections by Gram-negative pathogens (Fig 5E). The analysis suggests that primed neutrophils have metabolic and gene expression alterations linked to senescence, and also activation of innate immune signalling via MAPK signalling pathway to express pro-inflammatory cytokines, chemokines, and antimicrobial peptides. In addition, although most promoter regions of pro-inflammatory cytokines are not marked by H3K4me3, deposition of this mark was found in promoter regions of cytokine receptors *il10rb, tgfbr1b, ifngr1l, cxcr3.2* and *cxcr3.3* (Table S3). These results highlight the signalling pathways that may be activated and their importance for cell survival and immunomodulation.

### Reprogramed neutrophils produce more mitochondrial ROS for enhanced antimicrobial function

The ChIP-seq dataset strongly suggested increased mitochondrial ROS (mtROS) production upon *Shigella* WT priming (Fig 5E). In agreement, we found the promoter regions of the genes encoding MCU and VDAC transporters (mitochondrial permeability transition pore (mPTP)), as well as several genes involved in the tricarboxylic acid cycle and oxidative phosphorylation (Fig 6A and Table S3) are marked by H3K4me3 in *Shigella* WT-primed but not in naïve larvae. To test if mtROS production was increased in primed neutrophils, we used flow cytometry to quantify MitoTracker CM-H2XRos dye in naïve and *Shigella*-primed neutrophils in resting and re-stimulated state at 6 hp2i. Quantifications show that naïve neutrophils when infected with *Shigella* have reduced mtROS, compared to unstimulated neutrophils (Fig 6B). Primed neutrophils also present lower levels of mtROS when unstimulated, however after 6 hp2i mtROS levels are significantly increased as compared to unstimulated naïve and primed neutrophils.

**Figure 6.**
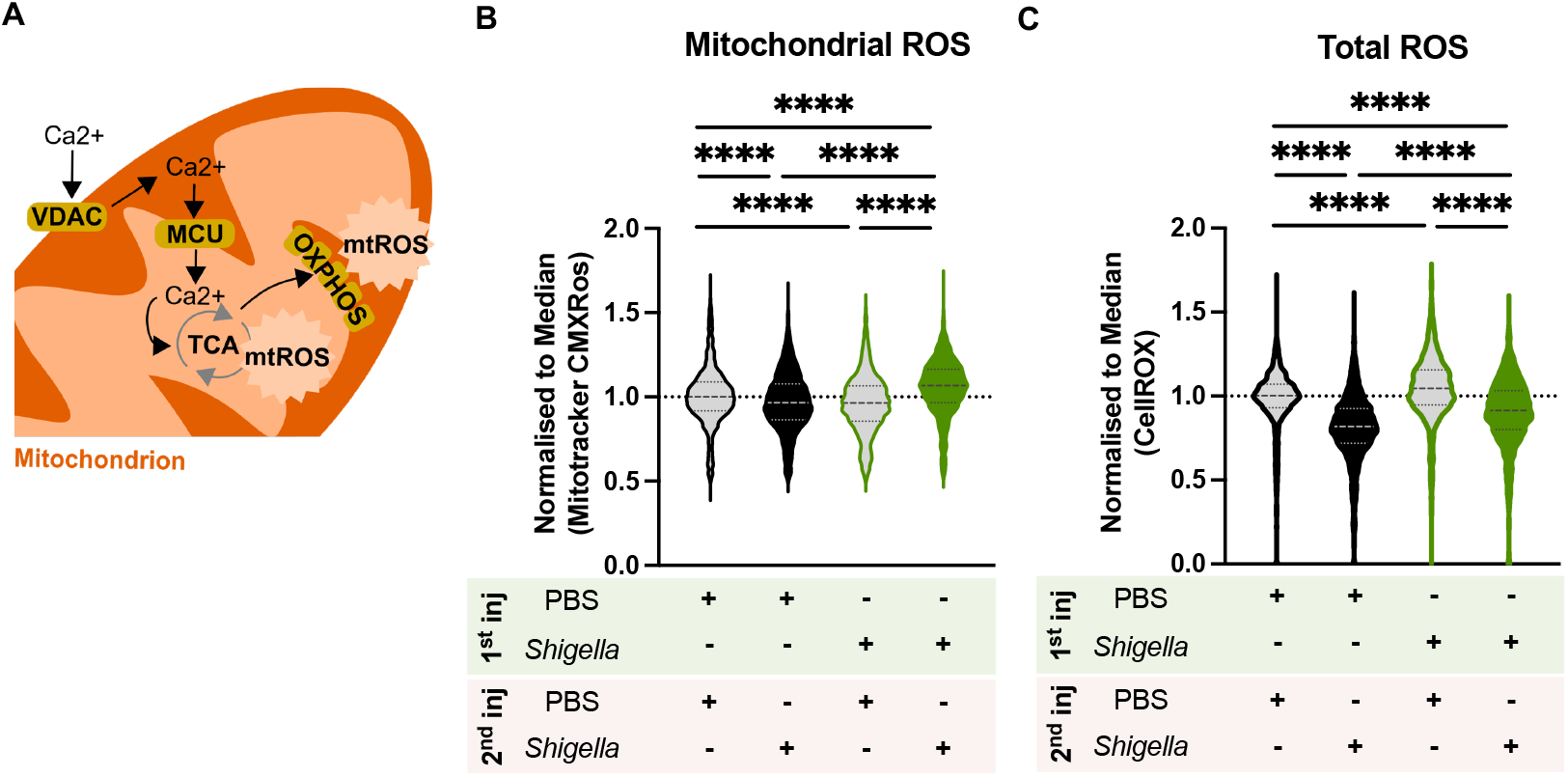
Trained neutrophils produce more ROS. **A** Increased calcium transportation into the mitochondrial inner membrane via the mitochondrial permeability transition pore (VDAC and MCU) increases mtROS produced by TCA and oxidative phosphorylation. **B** Normalised fluorescence of MitoTracker Red CM-H_2_XRos (red) dye in neutrophils from naïve (black outlines) and *Shigella*-primed (green outlines, 1.2×10^3^ ± 5.3×10^2^ CFUs) larvae (Tg(*mpx*::GFP)) unstimulated (grey fills) or infected with a lethal dose of *S. flexneri* M90T (PBS – black fill, 2.6×10^4^ ± 9.3×10^3^ CFUs, *Shigella* – green fill, 3×10^4^ ± 6.6×10^3^ CFUs) at 6 hp2i. Data pooled from 3 experiments with n>10 larvae per condition per experiment (mean ± SD, ****p < 0.0001, 2-way ANOVA with Sidak’s multiple comparisons test). **C** Normalised fluorescence of CellROX Deep Red dye in neutrophils from naïve (black outlines) and *Shigella*-primed (green outlines, 1.4×10^3^ ± 6.7×10^2^ CFUs) larvae (Tg(*mpx*::GFP)) unstimulated (grey fills) or infected with a lethal dose of *S. flexneri* M90T (PBS – black fill, 2.6×10^4^ ± 1×10^4^ CFUs, *Shigella* – green fill, 2.6×10^4^ ± 1×10^4^ CFUs) at 6 hp2i. Data pooled from 3 experiments with n>10 larvae per condition per experiment (mean ± SD, ****p < 0.0001, 2-way ANOVA with Sidak’s multiple comparisons test).

Given the redox crosstalk between mitochondria and NADPH oxidases ^61^, we investigated total ROS production in neutrophils using CellROX dye. Quantifications showed that naïve neutrophils when infected (6 hp2i) have significantly lower total ROS levels as compared to unstimulated state (Fig 6C). *Shigella*-primed neutrophils in resting phase showed higher levels of ROS, compared to naïve unstimulated neutrophils. However, when primed neutrophils became stimulated, total ROS levels decreased, but not to the same extent as in naïve infected neutrophils. The effect of both mtROS and ROS on bacterial killing was tested by adding specific inhibitors of mPTP (cyclosporin A (CsA)) and NADPH oxidase (diphenyleneiodonium chloride (DPI)) to naïve and *Shigella*-primed larvae bath during the 6 hp2i. Recovered CFU showed that inhibitors had more impact in bacterial burden control in *Shigella*-primed larvae than in naïve larvae (Fig S6B-C and S6E-F), suggesting that both mtROS and ROS are responsible for improved bacterial clearance in primed larvae.

To test if trained immunity can impact other aspects of host defence, we investigated the levels of myeloperoxidase (MPO) in trained neutrophils. Using the transgenic zebrafish line Tg(*mpx*::GFP) in which GFP expression is driven by the myeloperoxidase promoter, we quantified MPO production using FACS. In agreement with its absence from ChIP-seq analysis, we did not observe significant differences in MPO production between naïve and primed neutrophils when reinfected (Fig S6G), highlighting the specificity of the *Shigella* training towards mitochondrial ROS production.

Together these findings suggest that *Shigella*-trained neutrophils sustain H3K4me3 modifications that result in enhanced antimicrobial function via mtROS in preparation for a second challenge.

## DISCUSSION

Here we introduce the use of zebrafish larvae to study innate immune training, demonstrating that local infection by *Shigella*, BCG and β-glucan induce non-specific protective effects. The zebrafish larval model is a highly attractive training model – as compared to *in vitro* (single cell type system) and mouse models – as it is a vertebrate model with only innate immune responses during its first ~4 weeks of development.

Most work on trained immunity has been largely focused on monocytes and macrophages. However, there is growing evidence that neutrophils also play important roles upon training ^17,18^. In the case of *Shigella* infection, macrophages are unable to control infection as bacteria induce cell death programmes ^43,45,62^. Although neutrophils can also be killed by *Shigella*, they resist for longer than macrophages and can rapidly eliminate bacteria ^43,45^. Here we show that *Shigella* infection induces production of new neutrophils that are more protective and have better killing capacity than naïve neutrophils. We show that this enhanced infection control is linked to host control of its inflammatory state, which also contributes to increased survival upon a secondary infectious challenge. *Shigella*-priming protects against Gram-negative bacteria (as *Shigella* and *P. aeruginosa*) and also Grampositive bacteria (as *S. aureus*), suggesting that it induces heterologous immunological memory.

To test the hypothesis that *Shigella* induces long-lasting training mechanisms in zebrafish larvae, we extended the interval between infections to 5 days and observed protection against *Shigella* in those larvae. Although protection levels might wane, this is in agreement with the decline of BCG-induced protection after vaccination ^63,64^. In the future it will be of great interest to test adult zebrafish (> 3 months) as well as investigate if *Shigella*-training mechanisms can be transmitted trans-generationally in zebrafish as shown using mice ^65–67^. We envision that with the practices of zebrafish laboratories to maintain adequate levels of heterozygosity ^68^, by consistently outbreeding their stocks (unlike mouse lines) the zebrafish model can be used to complement the mouse model aiming to elucidate transmission of trained immunity in humans.

Work in zebrafish has shown that BCG vaccination protects against *Mycobacterium marinum* and that β-glucan (systemic) protects against *Salmonella* Typhimurium ^36,39^. Both BCG and β-glucan induced protection against *Shigella* infection, and yet had important differences that highlight distinct pathways that each stimuli uses to induce training (Table 1). Consistent with this, work using mice has shown that BCG induces training via NOD2 receptor and β-glucan via dectin-1 receptor ^52,53,69^. Although *Shigella*-priming phenotypes in zebrafish share similarities with those from BCG- and β-glucan-priming (such as non-specific protection and enhanced survival upon new infection), the H3K4me3 enriched pathways following *Shigella* training differ from those found in BCG-trained human neutrophils ^17^, which supports our conclusion that both stimuli induce different training mechanisms in neutrophils. Our findings suggest that *Shigella*-priming is multifactorial, depending on *Shigella* recognition by innate immune receptors, strong cytokine production, and an active T3SS.

Enriched pathways from our ChIP-seq datasets indicate that mtROS is playing an important role in enhancing antimicrobial activity of *Shigella*-primed neutrophils. Historically, ROS production in neutrophils has been associated with NADPH oxidase, and therefore the bactericidal role of mtROS in neutrophils has remained mostly elusive. However, recent work in human neutrophils suggested that mitochondria generated ROS enhanced bactericidal activity ^70^. In agreement, our work highlights that mtROS in neutrophils plays a key role in bacterial killing, and we envision that future therapeutic strategies could manipulate mtROS by using pro-oxidants (such as SkQN ^71^ or MitoK3 ^72^) specifically targeted to mitochondria in neutrophils.

Decades of research have tried to generate a vaccine with sufficient efficacy that could be used against different *Shigella* serotypes in humans ^73–75^. Our long-term goal is to guide vaccine studies in humans, exploiting innate immune memory to combat *Shigella* infections. A variety of animal models have significantly contributed to understanding shigellosis and feasibility of vaccines to combat it. We highlight zebrafish as an important animal model to investigate how innate immune training can be manipulated to control *Shigella* infection. It is next of great interest to train neutrophils using live *Shigella* in humans. In this way, we can discover fundamental components of immunity shared between zebrafish and humans that can be trained to promote a better immune response and combat *Shigella* infection.

## ACKNOWLEDGEMENTS

We thank Mostowy lab members, in particular Vincenzo Torraca, Damián Lobato-Márquez and Ana Teresa López-Jiménez, for helpful discussions. We thank Alain Filloux for providing *P. aeruginosa*. We thank Tracy Palmer for providing *S. aureus*. We thank Marcel Behr for providing BCG. We thank Felix Randow for providing *S. flexneri* Ruby strains. We thank the LSHTM Biological Services Facility for the work and care of our fish stocks. We thank Tomek Prasjnar for troubleshooting embryo dissociation protocols. We thank Imperial College London Flow Cytometry Facility South Kensington, in particular Larissa Zarate-Garcia and Jane Srivastava, for their help with neutrophil sorting. We thank Diagenode for the work and helpful discussions on ChIPseq. Research in the F.C.W. laboratory is supported by the BBSRC (BB/R016283/). Research in S.M. laboratory is supported by a European Research Council Consolidator Grant (772853 - ENTRAPMENT), Wellcome Trust Senior Research Fellowship (206444/Z/17/Z) and the Lister Institute of Preventive Medicine.

## AUTHOR CONTRIBUTIONS

Conceptualization M.C.G., S.M.

Methodology M.C.G., D.B., M.K.B.

Software M.C.G., D.B., M.K.B., F.C.W.

Validation M.C.G.

Formal analyses M.C.G., D.B., F.C.W.

Investigation M.C.G., D.B., M.K.B.

Resources

Data curation M.C.G.

Writing M.C.G., S.M.

Visualization

Supervision S.M.

Project administration S.M.

Funding acquisition S.M.

## MATERIALS AND METHODS

### Ethics statements

Animal experiments were performed according to the Animals (Scientific Procedures) Act 1986 and approved by the Home Office (Project licenses: PPL P84A89400 and P4E664E3C).

### Zebrafish husbandry

Embryos were obtained from naturally spawning zebrafish and larvae were maintained at 28.5°C in embryo medium (0.5X E2 medium), unless specified otherwise. Transgenic zebrafish lines used here were Tg(*lyz*::dsRed)^nz50 76^, Tg(*mpx*::GFP)^1114 77^, Tg(*mpeg1*::Gal4-FF)^gl25^/Tg(UAS-E1b::*nfsB*.mCherry)^c264 78^. For injections and live microscopy, larvae were anaesthetized with 200 μg/ml tricaine (Sigma-Aldrich) in embryo medium. For injections in larvae > 5 dpf, lidocaine (2 μg/L) was used as an analgesic and added to the embryo medium (together with 200 μg/ml tricaine (Sigma-Aldrich)) prior to injection. After checking for full recovery from the anaesthetic, larvae were kept in E2 medium with lidocaine (2 μg/L) for 18-24 h at 28.5°C. Larvae were monitored for appearance, righting reflex, reactive reflex and opercular movement from day 5 dpf for 2-3 times a day. Larvae were not fed during the course of the experiment ^79^.

### Zebrafish injections

Bacterial strains used in this study can be found in Table S4. *Shigella, Pseudomonas* and *Staphylococcus* strains were grown on trypticase soy agar (TSA, Sigma-Aldrich) plates with the appropriate antibiotics – for *Shigella* growth, plates were supplemented with 0.01% Congo red (Sigma-Aldrich). Overnight cultures were grown from individual colonies at 37°C and 200 rpm, in 5 ml trypticase soy broth (TSB, Sigma-Aldrich and Oxoid for *Staphylococcus*) supplemented with the appropriate antibiotics as above. For injections, bacteria were grown until an optical density (OD) of 0.55–0.65 at 600 nm (log phase) by diluting the overnight culture 50X in fresh TSB supplemented with the appropriate antibiotics. When in log phase, bacteria were spun down and washed in phosphate buffer saline (PBS, Sigma-Aldrich). The desired concentration was achieved by resuspension of bacteria in injection buffer (2% polyvinyl-pyrrolidone (PVP, Sigma-Aldrich) in PBS and 0.5% phenol red (Sigma-Aldrich)). Control groups were injected with injection buffer (referred as PBS/naïve group).

For inactivated bacteria injections suspensions were prepared as described above and, prior to injection, were incubated for 30 min at 60°C for heat-killing or incubated in 4% paraformaldehyde (PFA, Sigma-Aldrich) for 30 min at RT for PFA-killing.

BCG was grown in 7H9 broth (supplemented with 10 % ADC enrichment medium, 0.2 % glycerol and 0.02 % Tween-80) until OD 0.5-0.6. The culture was then spun down and washed with PBS. The suspension was then centrifuged and resuspended in PVP to prevent large clumps. Single cell suspensions were obtained by passing the bacteria through different gauge syringes (25G, 27G and 29G) 10 times each. After achieving the desired concentration, bacteria were resuspended in injection buffer with higher concentration of PVP (4%).

A stock solution of β-glucan (Invivo Biosystems cat no. tlrl-bgp) was prepared at 20mg/mL in PBS and stored at 4 °C for a maximum of a month. For injections, indicated dilutions were prepared in injection buffer. Different amounts of β-glucan were first injected in the HBV of larvae to test for dose dependent protection (Fig S3C-E). The highest amount tested (20 μg/μL) was chosen as it suggested stronger protection responses.

Lipopolysaccharide (LPS, Sigma-Aldrich cat no. L6143) was prepared to mimic the amount of bacteria injected in a non-lethal dose (2500 CFU). According to ^80^, 2500 bacterial cells in exponential phase would correspond to 0.125 ng of LPS. Therefore, a stock solution of 0.25 mg/mL LPS in PBS was prepared, and prior before injection was mixed 1:1 with 0.5% phenol red.

For injection, 1–2 nl of bacterial / β-glucan / LPS suspension or control solution were microinjected in the hindbrain ventricle (HBV) of zebrafish larvae.

For bacterial enumeration, larvae tissues were disrupted in 0.4% Triton X-100 (Sigma-Aldrich) with the aid of mechanical pestles, and serial dilutions were plated in the appropriate medium at the indicated timepoints. Plates were incubated at 37°C and colony forming units (CFU) were enumerated when colonies were visible.

### Design of *S. flexneri ΔospF* mutant

*S. flexneri ΔospF* mutant was created using a λ-Red-mediated recombination ^81^. In brief, pKD4 plasmid was used as template to amplify a kanamycin resistance-encoding DNA cassette using primers containing 50 bp nucleotides homologous to the site of insertion (Table S4). *S. flexneri* electrocompetent cells producing λ-Red recombinase were electroporated with resulting PCR fragments. Electroporated cells were then plated in TSA plates supplemented with 0.01% of Congo red and 50□μg/ml of kanamycin. To remove the kanamycin resistance-encoding DNA cassette, bacteria were transformed with pCP20 plasmid (that encodes the yeast *flp* recombinase, ^82^). Deletions were verified by PCR using confirmation primers in Table S4.

### Microscopy and image analysis

Whole larvae and HBV images were acquired using stereo fluorescent microscope Leica M205FA (Leica, Germany), and by placing anaesthetised larvae on 1% agarose-E2 plates.

Image files were processed using ImageJ/FIJI software. Whole larvae leukocyte quantification was performed according to ^83^ with modifications. In brief, fluorescent images were converted to binary in ImageJ/FIJI, resulting in images in which fluorescence was converted into black pixels onto a white background. For each image, pixels of 5 individual cells were quantified. Total pixel count of each larvae was divided by the average of 5 individual leukocytes to determine total number of cells. Leukocyte quantification in the HBV was determined manually in the region highlighted in Fig 1G.

### Zebrafish chemical treatments

For inflammasome stimulation, larvae were kept at 28.5°C in E2 medium with 0.1 μM Nigericin (Sigma-Aldrich cat no. 481990-5MG) or 1 % D-glucose (Sigma-Aldrich cat no. G8270-100G) ^28^ for 24, 48 or 96 h from 2 dpf. For NAPDH ROS and mitochondrial ROS inhibition after the 2^nd^ infection, larvae were kept at 28.5°C in E2 medium with 100 μM diphenyleneiodonium chloride (DPI, Sigma-Aldrich cat no. D2926) ^84^ and 10 μM cyclosporin A (CsA, Enzo Life Sciences cat no. BML-A195-0100) ^85^, both in DMSO, for 6 h until larvae were dissociated for flow cytometry. Control larvae were kept in E2 with 1% DMSO.

### RNA extraction, cDNA synthesis and qRT-PCR

RNA was extracted from 5-10 snap-frozen larvae with the RNeasy Mini kit (Qiagen cat no. 74104) and reverse-transcribed using QuantiTect Reverse Transcription kit (Qiagen cat no 205311) according to manufacturer’s instructions. Quantitative PCR (qPCR) was performed using 7500 Fast Real-Time PCR System machine and 7500 Fast Real-Time PCR software v2.3 (Applied Biosystems, Foster City, California) and SYBR green master mix (Applied Biosystems cat no. 10187094). Template cDNA was subjected to PCR using primers described in Table S4 and samples were run in technical duplicates. The comparative Ct method was used for gene expression quantification, and *ef1a1l1* was used as the housekeeping gene.

### Flow cytometry

Tg(*lyz*::dsRed)^nz50^ and Tg(*mpx*::GFP)^i114^ transgenic larvae were dissociated for flow cytometry following the protocol from ^86^, with some modifications. In brief, 20 to 100 larvae were washed twice in calcium-free Ringer’s solution for 10 min with slow shaking at RT. After removal of Ringer’s solution, 2 mL of dissociation solution (0.5% Trypsin EDTA 10x no phenol (Fisher cat no.15400054)) was added. Larvae were incubated twice for 10 min at 28.5°C, with gentle pipetting in between incubations to help tissue digestion. The reaction was stopped by adding 10% foetal calf serum (Fisher cat no. 11550356). The suspensions were sieved using 70 μm Nylon cell strainers (Merck CLS431751-50EA) or cell strainer snap caps from test tubes (Corning 352235) and then gently centrifuged for 3 min at 800g. Cells were then washed with 1mL PBS with 10 mM HEPES and 1 mM EDTA. For live/dead staining, cells were incubated with either LIVE/DEAD™ fixable near IR dead cell stain (1:2000, Thermofisher cat no. L34994) or LIVE/DEAD™ fixable violet dead cell stain (1:2000, Thermofisher cat no. L34964) dyes for 30 min at 4°C. Cells were fixed with 4% PFA for 10 min at RT for flow cytometry. For ROS staining, 5μM CellROX™ Deep Red (Thermofisher cat no. C10422) was added to the E2 medium for 30 min prior to larvae dissociation and the larvae were kept in the dark at 28.5°C. Single cells were measured on a LSRII (BD Biosciences). For mitochondrial ROS staining, 300 nM MitoTracker™ Red CM-H_2_XRos (Thermofisher cat no. M7513) was added to the dissociated cells for 30 min at 28.5°C. Single cells were measured on a FACSAria III cell sorter (BD Biosciences). Data was analysed with FlowJo software v10.7.1. To prepare samples for ChIP-seq, cells were fixed with 1% formaldehyde for 8 min at RT with occasional mixing. To stop the reaction 100 μL of glycine 1.25 M were added and the samples were incubated for 5 min at RT. After centrifuging at 600g for 10 min at 4°C, cells were washed with ice cold PBS containing protease inhibitor cocktail (Sigma-Aldrich cat no. 5892970001) and centrifuged again. Cells were then resuspended in PBS with 10mM HEPES and 1 mM EDTA. FACS sorting was done on a FACSAria III cell sorter (BD Biosciences). Data was analysed with BD FACSDiva software. Gating strategies can be found in Fig S5A for neutrophil sorting, in Fig S6A for mitochondrial ROS measurement and Fig S6D for CellROX measurement.

### ChIP-seq and analysis

For ChIP experiments, a minimum of 20 000 sorted Tg(*<u>lyz</u>*:dsRed) neutrophils were collected per sample (PBS, *Shigella, Shigella* ΔT3SS) and per replicate (each biological replicate corresponds to two experiments). Cells were kept at −80oC until sequencing. ChIP-seq was conducted by Diagenode ChIP-seq/ChIP-qPCR Profiling service (Diagenode cat no. G02010000). Chromatin was prepared for ChIPseq using the True MicroChIP Kit (Diagenode cat no. C01010130) and was sheared using Bioruptor® Pico sonication device (Diagenode cat no. B01060001) combined with the Bioruptor® Water cooler for 5 cycles using a 30’’ [ON] 30’’ [OFF] settings. Optimization of shearing conditions was done on chromatin corresponding to 10 000 cells. Immunoprecipitation was done on chromatin corresponding to the remaining 10 000 cells using antibodies against H3K4me3 (Diagenode, cat no. C15410003) – 10% of the chromatin was used for input. Libraries were prepared using IP-Star® Compact Automated System (Diagenode cat no. B03000002) from input and ChIP’d DNA using MicroPlex Library Preparation Kit v2 (12 indices) (Diagenode cat no. C05010013). Libraries were pooled and sequenced with Illumina Novaseq 6000, running NovaSeq Control Software 1.6.0, with paired-end reads of 50bp length. Quality control of sequencing reads was performed using FastQC ^87^.

ChIP-sequencing reads were analysed using the Galaxy platform (usegalaxy.eu). Reads were aligned to the reference genome GRCz10 using the Bowtie2 software ^88^ and PCR duplicates were removed with MarkDuplicates ^89^. Peak calling was performed using MACS2 callpeak ^90^ and the narrow peaks lists were used in IDR ^91^ to identify reproducible peaks between replicates. Finally, ChIPseeker ^92^ was used for peak annotation with a promoter defined as ±3kb around the transcription start site. IGV v2.9.4 ^93^ was used for peak visualisation.

For pathway annotation, the lists of genes associated with promoter regions highlighted in bold in Fig 5B and 5D, Fig S5B and S5D (PBS, *Shigella* WT, *Shigella* ΔT3SS) were analysed against KEGG pathways in Cytoscape v3.9.1 ^94^ using the plug-in ClueGO v2.5.9 ^95^.

### Resource availability

#### Lead Contact

Further information and requests for resources and reagents should be directed to and will be fulfilled by the Lead Contact, Serge Mostowy at London School of Hygiene & Tropical Medicine, United Kingdom.

#### Materials Availability

All unique/stable reagents generated in this study are available from the Lead Contact without restriction.

#### Data and Code Availability

The ChIP sequencing data of neutrophils 48 h post *Shigella* priming as reported in this paper can been found in the NCBI Gene Expression Omnibus under accession number GEO: GSE217063.

### Statistical analysis

Statistical significance was determined using GraphPad Prism v9. For differences in survival curves, the Log-rank (Mantel-Cox) test was used. Data from bacterial burden and gene expression levels were Log10- or Log2-transformed, respectively. Pairwise comparisons were determined using a Student’s unpaired *t*-test. For multiple comparisons, one-way or two-way ANOVA tests with Sidak’s correction were used, as indicated in the figure legend.

## SUPPLEMENTAL TABLES

**Table S1, related to Figure 5 – Gene annotation from ChIP-seq analysis of PBS treated larvae.**

**Table S2, related to Figure 5 – Gene annotation from ChIP-seq analysis of *Shigella* ΔT3SS-primed neutrophils**

**Table S3, related to Figure 5 – Gene annotation from ChIP-seq analysis of *Shigella*-primed neutrophils**

**Table S4.**
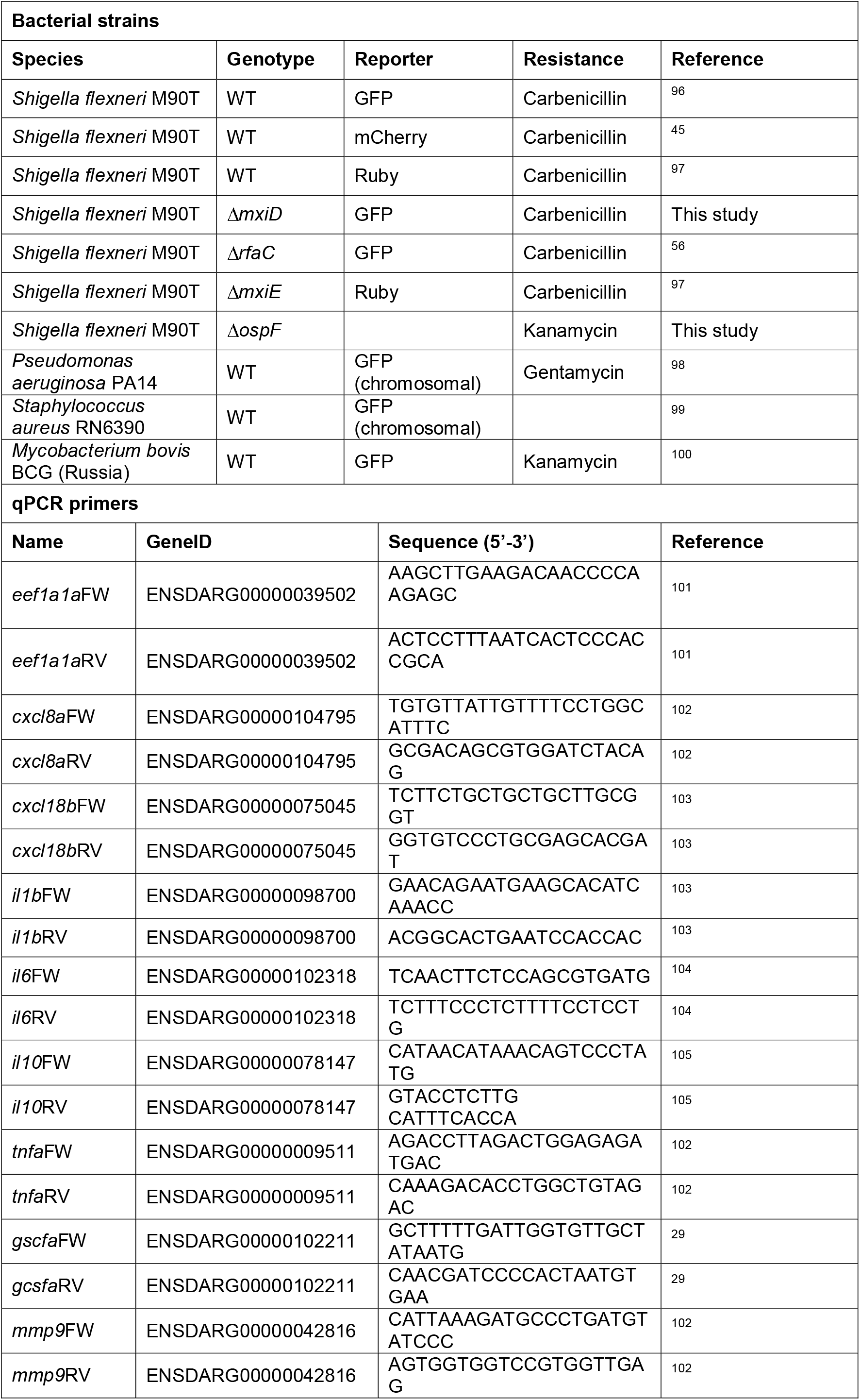
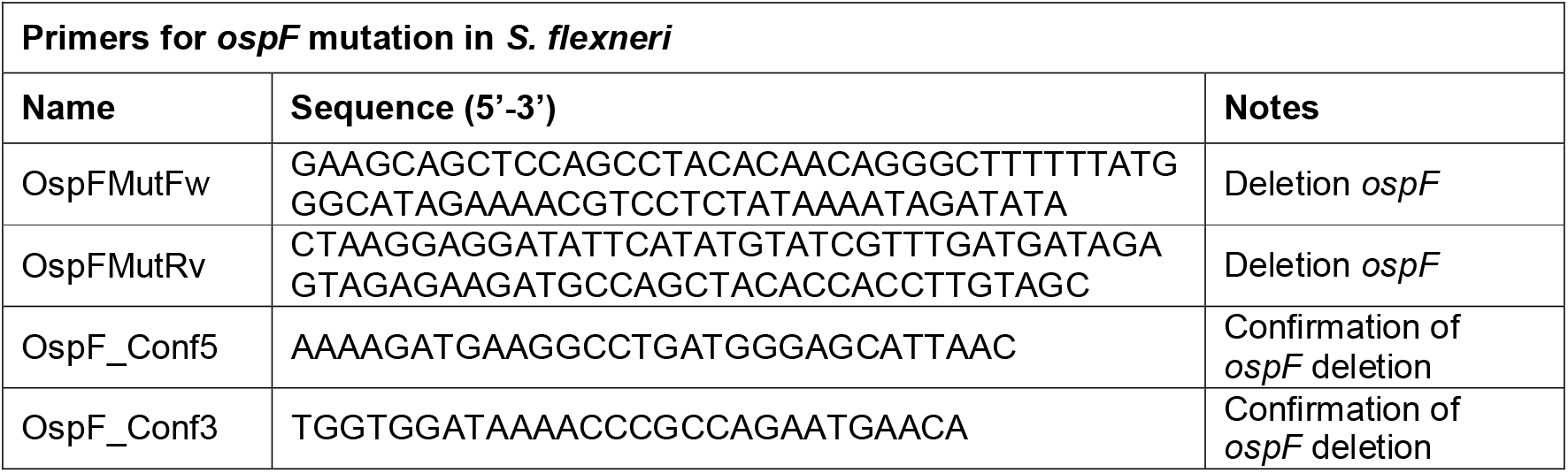
Bacterial strains, qPCR primers and primers for *ospF* mutation in *S. flexneri* used in this study.

## SUPPLEMENTAL FIGURES

**Figure S1, related to Figure 1 – Expression of *il6* and *tnfa* during priming and reinfection.**

**A** Fold change in the expression of *il6* and *tnfa* in larvae injected with non-lethal dose of *S. flexneri* M90T (green full bars, 1.3×10^3^ ± 5.5×10^2^ CFUs) as compared to PBS control (naïve-black full bars). Data pooled from 3 experiments with n>5 larvae per time point per condition per experiment (mean ± SD, *p < 0.05, 2-way ANOVA with Sidak’s multiple comparisons test).

**B** Log_10_-transformed CFU counts *Shigella*-primed larvae injected with a non-lethal dose of *S. flexneri* M90T (1.3×10^3^ ± 5.5×10^2^ CFUs). Data pooled from 3 independent experiments using n=3 larvae per condition per experiment (mean ± SD, ***p < 0.001, ****p < 0.0001, 1-way ANOVA with Dunnett’s multiple comparisons test).

**C** Fold change in the expression of *il6* and *tnfa* following lethal dose injection of *S. flexneri* M90T in naïve (black full bars, 2.5×10^4^ ± 1×10^4^ CFUs) and *Shigella*-primed (green full bars, 2.7×10^4^ ± 4.5×10^3^ CFUs) as compared to PBS injected controls (grey bars). Data pooled from 3 experiments with n>5 larvae per time point per condition per experiment (mean ± SD, **p < 0.01, 2-way ANOVA with Sidak’s multiple comparisons test).

**Figure S2, related to Figure 2 – Bacterial burden during reinfection with *P. aeruginosa* and *S. aureus*.**

**A** Log_10_-transformed CFU counts of naïve and *Shigella*-primed (in green section: 2.2×10^3^ ± 3.8×10^2^ CFUs) larvae injected with a lethal dose of *P. aeruginosa* (in orange section: PBS – 2.6×10^3^ ± 7.4×10^2^ CFUs, *Shigella* – 2.3×10^3^ ± 5.5×10^2^ CFUs). Data pooled from 3 independent experiments using n=3 larvae per condition per experiment (mean ± SD, ****p < 0.0001, 2-way ANOVA with Dunnett’s multiple comparisons test).

**B** Log_10_-transformed CFU counts of naïve and *Shigella*-primed (in green section: 1.8×10^3^ ± 7×10^2^ CFUs) larvae injected with a non-lethal dose of *S. aureus* (PBS – 1.7×10^4^ ± 4.5×10^3^ CFUs, *Shigella* – 2.6×10^4^ ± 1×10^4^ CFUs). Data pooled from 3 independent experiments using n=3 larvae per condition per experiment (mean ± SD, ***p < 0.001, ****p < 0.0001, 1-way ANOVA with Dunnett’s multiple comparisons test).

**Figure S3, related to Figure 3 – Leukocyte recruitment to BCG infected HBV, testing different β-glucan concentrations, and granulopoiesis in β-glucan priming.**

**A** Quantification of recruited neutrophils (Tg(*lyz*::dsRed)) to the HBV in naïve (black full circles) and BCG-primed (purple full circles, 4×10^2^ ± 1.5×10^2^ CFUs) larvae. Data pooled from 3 experiments with n>4 larvae per time point per condition per experiment (mean ± SD, 2-way ANOVA with Sidak’s multiple comparisons test).

**B** Quantification of recruited macrophages (Tg(mpeg::mCherry)) to the HBV in naïve (black full circles) and BCG-primed (purple full circles, 4×10^2^ ± 1.5×10^2^ CFUs) larvae. Data pooled from 3 experiments with n>4 larvae per time point per condition per experiment (mean ± SD, 2-way ANOVA with Sidak’s multiple comparisons test).

**C** Survival curves from naïve (black full circles) and β-glucan-primed (light blue full circles: 5 ng, dark blue full circles: 10 ng) larvae infected with a lethal dose of *S. flexneri* M90T. Data pooled from 3 independent experiments using n>18 larvae per condition per experiment (**p < 0.01, Log-rank (Mantel-Cox) test).

**D** Log_10_-transformed CFU counts of naïve and β-glucan-primed larvae injected with a lethal dose of *S. flexneri* M90T (PBS – 2.5×10^4^ ± 7.3×10^3^ CFUs, β-glucan 5 ng – 2.7×10^4^ ± 3.2×10^3^ CFUs, β-glucan 10 ng– 2.7×10^4^ ± 5.1×10^3^ CFUs). Data pooled from 3 independent experiments using n=3 larvae per condition per experiment (mean ± SD, 2-way ANOVA with Sidak’s multiple comparisons test).

**E** Quantification of neutrophils (Tg(*lyz*::dsRed)) at 0, 24 and 48 hp1i injected with PBS (naïve-black full circles) and a non-lethal dose of β-glucan (light blue full circles: 5 ng, dark blue full circles: 10 ng). Data pooled from 3 experiments with n>8 larvae per time point per condition per experiment (mean ± SD, *p < 0.05, 2-way ANOVA with Sidak’s multiple comparisons test).

**F** Fold change in the expression of *gcsfa* following non-lethal dose injection of β-glucan (blue bars, 20 ng) as compared to PBS injected controls (black bars). Data pooled from 3 experiments with n=10 larvae per time point per condition per experiment (mean ± SD, **p < 0.01, unpaired Student’s *t*-test).

**G** Quantification of neutrophils (Tg(*lyz*::dsRed)) in the AGM at 48 hp1i injected with PBS (naïve-black full circles) and a non-lethal dose of β-glucan (blue full circles, 20 ng). Data pooled from 3 experiments with n>8 larvae per time point per condition per experiment (mean ± SD, unpaired Student’s *t*-test).

**Figure S4, related to Figure 4 – Bacterial burden during reinfection upon stimulation of inflammatory pathways and virulence assays with *Shigella* mutants.**

**A** Log_10_-transformed CFU counts from E2 (embryo media control, black full circles) and nigericin (blue full circles, 0.1 μM) treated larvae infected with a lethal dose of *S. flexneri* M90T (PBS – 2.7×10^4^ ± 8.5×10^3^ CFUs, nigericin – 2.6×10^4^ ± 4.1×10^3^ CFUs). Data pooled from 3 independent experiments using n=3 larvae per condition per experiment (mean ± SD, 2-way ANOVA with Sidak’s multiple comparisons test).

**B** Log_10_-transformed CFU counts from E2 (embryo media control, black full circles) and glucose (blue full circles, 1%) treated larvae infected with a lethal dose of *S. flexneri* M90T (PBS – 2.7×10^4^ ± 8.5×10^3^ CFUs, glucose – 2.7×10^4^ ± 1×10^4^ CFUs). Data pooled from 3 independent experiments using n=3 larvae per condition per experiment (mean ± SD, 2-way ANOVA with Sidak’s multiple comparisons test).

**C** Log_10_-transformed CFU counts from naïve (black full circles), live *Shigella*-primed (green full circles, 1.3×10^3^ ± 6.6×10^2^ CFUs), heat-killed (HK) *Shigella*-primed (dark green/white squares) and PFA-killed *Shigella*-primed (dark green/white triangles) larvae infected with a lethal dose of *S. flexneri* M90T (PBS – 2.7×10^4^ ± 6.1×10^3^ CFUs, *Shigella* – 2.7×10^4^ ± 1×10^4^ CFUs, *Shigella* HK – 3.2×10^4^ ± 3.3×10^4^ CFUs, *Shigella* PFA – 3.2×10^4^ ± 1.7×10^4^ CFUs). Data pooled from 3 independent experiments using n=3 larvae per condition per experiment (mean ± SD, **p < 0.01, ****p < 0.0001, 2-way ANOVA with Sidak’s multiple comparisons test).

**D** Log_10_-transformed CFU counts from naïve (black full circles), *Shigella*-primed (green full circles, 1.6×10^3^ ± 6.5×10^2^ CFUs) and LPS-primed (moss green full circles, 0.125 ng) larvae infected with a lethal dose of *S. flexneri* M90T (PBS – 1.9×10^4^ ± 6.3×10^3^ CFUs, *Shigella* – 2.8×10^4^ ± 4.7×10^3^ CFUs, LPS – 1.9×10^4^ ± 9.1×10^3^ CFUs). Data pooled from 3 independent experiments using n=3 larvae per condition per experiment (mean ± SD, 2-way ANOVA with Sidak’s multiple comparisons test).

**E** Survival curves from naïve (black full circles), *Shigella*-primed (green full circles, 1.7×10^3^ ± 5.5×10^2^ CFUs) and LPS-primed (moss green full circles, 0.5 ng) larvae infected with a lethal dose of *S. flexneri* M90T (PBS – 2×10^4^ ± 4.6×10^3^ CFUs, *Shigella* – 2.7×10^4^ ± 8.2×10^3^ CFUs, LPS – 2.3×10^4^ ± 1.2×10^4^ CFUs). Data pooled from 3 independent experiments using n>8 larvae per condition per experiment. (****p < 0.0001, Log-rank (Mantel-Cox) test).

**F** Log_10_-transformed CFU counts from naïve (black full circles), *Shigella*-primed (green full circles, 1.7×10^3^ ± 5.5×10^2^ CFUs) and LPS-primed (moss green full circles, 0.5 ng) larvae infected with a lethal dose of *S. flexneri* M90T (PBS – 2×10^4^ ± 4.6×10^3^ CFUs, *Shigella* – 2.7×10^4^ ± 8.2×10^3^ CFUs, LPS – 2.3×10^4^ ± 1.2×10^4^ CFUs). Data pooled from 3 independent experiments using n=3 larvae per condition per experiment (mean ± SD, **p < 0.01, 2-way ANOVA with Sidak’s multiple comparisons test).

**G** Log_10_-transformed CFU counts from naïve (black full circles), *Shigella*-primed (green full circles, 1.6×10^3^ ± 6.5×10^2^ CFUs) and Shigella Δ*rfaC*-primed (moss green full circles, 8.5×10^2^ ± 7.5×10^2^ CFUs) larvae infected with a lethal dose of *S. flexneri* M90T (PBS – 1.9×10^4^ ± 6.3×10^3^ CFUs, *Shigella* – 2.8×10^4^ ± 4.7×10^3^ CFUs, Δ*rfaC* – 2.4×10^4^ ± 8.4×10^3^ CFUs). Data pooled from 3 independent experiments using n=3 larvae per condition per experiment (mean ± SD, **p < 0.01, 2-way ANOVA with Sidak’s multiple comparisons test).

**H** Survival curves from virulence assay of 3 dpf larvae infected with *Shigella* (green full circles, 9.6×10^3^ ± 2.5×10^3^ CFUs) and *Shigella ΔrfaC* (moss green full circles, 9.1×10^3^ ± 3.2×10^2^ CFUs). Data pooled from 3 independent experiments using n>8 larvae per condition per experiment. (****p < 0.0001, Log-rank (Mantel-Cox) test).

**I** Log_10_-transformed CFU counts from virulence assay of 3 dpf larvae infected with *Shigella* (green full circles, 9.6×10^3^ ± 2.5×10^3^ CFUs) and *Shigella ΔrfaC* (moss green full circles, 9.1×10^3^ ± 3.2×10^2^ CFUs). Data pooled from 3 independent experiments using n=3 larvae per condition per experiment (mean ± SD, 2-way ANOVA with Sidak’s multiple comparisons test).

**J** Fold change in the expression of *il1b, cxcl18b* and *mmp9* following injection of PBS (black full bars), *Shigella* (green full bars, 9.6×10^3^ ± 2.5×10^3^ CFUs) and *Shigella ΔrfaC* (moss green full bars, 9.1×10^3^ ± 3.2×10^2^ CFUs). Data pooled from 3 experiments with n>5 larvae per time point per condition per experiment (mean ± SEM, *p < 0.05, **p < 0.01, ***p < 0.001, 2-way ANOVA with Sidak’s multiple comparisons test).

**K** Log_10_-transformed CFU counts from naïve (black full circles), *Shigella*-primed (green full circles, 7.9×10^2^ ± 5.5×10^2^ CFUs) and Shigella Δ*mxiE*-primed (moss green full circles, 8.8×10^2^ ± 5.8×10^2^ CFUs) larvae infected with a lethal dose of *S. flexneri* M90T (PBS – 1.9×10^4^ ± 7.5×10^3^ CFUs, *Shigella* – 2.5×10^4^ ± 4.6×10^3^ CFUs, Δ*mxiE* – 2.2×10^4^ ± 7.6×10^3^ CFUs). Data pooled from 3 independent experiments using n=3 larvae per condition per experiment (mean ± SD, 2-way ANOVA with Sidak’s multiple comparisons test).

**L** Survival curves from naïve (black full circles), *Shigella*-primed (green full circles, 3.4×10^3^ ± 2.2×10^3^ CFUs) and *Shigella* Δ*ospF*-primed (moss green full circles, 2.9×10^3^ ± 1.1×10^3^ CFUs) larvae infected with a lethal dose of *S. flexneri* M90T (PBS – 2.4×10^4^ ± 8.5×10^3^ CFUs, *Shigella* – 2.7×10^4^ ± 1.5×10^3^ CFUs, Δ*ospF* – 2.8×10^4^ ± 8.9×10^3^ CFUs). Data pooled from 3 independent experiments using n>10 larvae per condition per experiment. (****p < 0.0001, Log-rank (Mantel-Cox) test).

**M** Log_10_-transformed CFU counts from naïve (black full circles), *Shigella*-primed (green full circles, 3.4×10^3^ ± 2.2×10^3^ CFUs) and *Shigella* Δ*ospF*-primed (moss green full circles, 2.9×10^3^ ± 1.1×10^3^ CFUs) larvae infected with a lethal dose of *S. flexneri* M90T (PBS – 2.4×10^4^ ± 8.5×10^3^ CFUs, *Shigella* – 2.7×10^4^ ± 1.5×10^3^ CFUs, Δ*ospF* – 2.8×10^4^ ± 8.9×10^3^ CFUs). Data pooled from 3 independent experiments using n=3 larvae per condition per experiment (mean ± SD, *p < 0.05, 2-way ANOVA with Sidak’s multiple comparisons test).

**N** Survival curves from virulence assay of 3 dpf larvae infected with *Shigella* (green full circles, 2.3×10^4^ ± 9.6×10^3^ CFUs) and *Shigella ΔmxiE* (moss green full circles, 2.2×10^4^ ± 9.8×10^3^ CFUs). Data pooled from 3 independent experiments using n=20 larvae per condition per experiment. (*p < 0.05, Log-rank (Mantel-Cox) test).

**O** Log_10_-transformed CFU counts from virulence assay of 3 dpf larvae infected with *Shigella* (green full circles, 2.3×10^4^ ± 9.6×10^3^ CFUs) and *Shigella ΔmxiE* (moss green full circles, 2.2×10^4^ ± 9.8×10^3^ CFUs). Data pooled from 3 independent experiments using n=3 larvae per condition per experiment (mean ± SD, 2-way ANOVA with Sidak’s multiple comparisons test).

**P** Survival curves from virulence assay of 3 dpf larvae infected with *Shigella* (green full circles, 2.3×10^4^ ± 1.1×10^3^ CFUs) and *Shigella ΔospF* (moss green full circles, 2.1×10^4^ ± 1.3×10^4^ CFUs). Data pooled from 2 independent experiments using n>15 larvae per condition per experiment. (*p < 0.05, Log-rank (Mantel-Cox) test).

**Q** Log_10_-transformed CFU counts from virulence assay of 3 dpf larvae infected with *Shigella* (green full circles, 2.3×10^4^ ± 1.1×10^3^ CFUs) and *Shigella ΔospF* (moss green full circles, 2.1×10^4^ ± 1.3×10^4^ CFUs). Data pooled from 2 independent experiments using n=3 larvae per condition per experiment (mean ± SD, 2-way ANOVA with Sidak’s multiple comparisons test).

**Figure S5, related to Figure 5 – ChIP-seq analysis of PBS, *Shigella*- and *Shigella* ΔT3SS-primed neutrophils.**

**A** Gating strategy for FACS sorting neutrophils from Tg(*lyz*::dsRed) larvae. Post cell identification, duplet exclusion and determination of live and dead cells (LIVE/DEAD™ Fixable Near IR Stain), neutrophils were identified by dsRed expression, followed by single cell sorting.

**B** Venn diagram showing number of genes marked by H3K4me3 peaks in their promoter regions (±3kb from TSS) of PBS treated larvae.

**C** Enriched KEGG-pathways (*p*-value < 0.01) associated with the 1049 genes marked by a H3K4me3 peak in their promoter regions (±3kb from TSS) in PBS treated larvae.

**D** Venn diagram showing number of common and unique genes marked by H3K4me3 peaks in their promoter regions (±3kb from TSS) of PBS, *Shigella* WT and *Shigella* ΔT3SS primed larvae.

**G** Enriched KEGG-pathways (*p*-value < 0.01) associated with the 854 genes marked by a H3K4me3 peak in their promoter regions (±3kb from TSS) common to *Shigella* WT and *Shigella* ΔT3SS primed larvae.

**Figure S6, related to Figure 6 – Quantifying ROS in primed neutrophils.**

**A** Gating strategy for flow cytometry of neutrophils from Tg(*mpx*::GFP) larvae. Neutrophils were identified by *mpx-gfp* expression. Neutrophils were analysed for their mitochondrial ROS production by fluorescence intensity detection of MitoTracker Red CM-H2Xros.

**B** Normalised fluorescence of MitoTracker Red CM-H_2_XRos dye neutrophils treated with CsA from naïve (black outlines) and *Shigella*-primed (green outlines, 1.2×10^3^ ± 5.3×10^2^ CFUs) larvae (Tg(*mpx*::GFP)) unstimulated (grey fills) or infected with a lethal dose of *S. flexneri* M90T (PBS – black fill, 2.6×10^4^ ± 9.3×10^3^ CFUs, *Shigella* – green fill, 3×10^4^ ± 6.6×10^3^ CFUs) at 6 hp2i. Data pooled from 3 experiments with n>10 larvae per condition per experiment (mean ± SD, ****p < 0.0001, 2-way ANOVA with Sidak’s multiple comparisons test).

**C** Log_10_-transformed CFU counts of naïve and *Shigella*-primed (1.2×10^3^ ± 5.3×10^2^ CFUs) larvae injected with a lethal dose of *S. flexneri* M90T (PBS – black fill, 2.6×10^4^ ± 9.3×10^3^ CFUs, *Shigella* – green fill, 3×10^4^ ± 6.6×10^3^ CFUs) and treated with CsA. Data pooled from 3 independent experiments using n=3 larvae per condition per experiment (mean ± SD, 2-way ANOVA with Sidak’s multiple comparisons test).

**D** Gating strategy for flow cytometry of neutrophils from Tg(*mpx*::GFP) larvae. Neutrophils were identified by *mpx-gfp* expression, followed by duplet exclusion and determination of live and dead cells (LIVE/DEAD™ Fixable Violet Stain). Live neutrophils were analysed for their total ROS production by fluorescence intensity detection of CellROX Deep Red.

**E** Normalised fluorescence of CellROX Deep Red dye neutrophils treated with DPI from naïve (black outlines) and *Shigella*-primed (green outlines, 1.4×10^3^ ± 6.7×10^2^ CFUs) larvae (Tg(*mpx*::GFP)) unstimulated (grey fills) or infected with a lethal dose of *S. flexneri* M90T (PBS – black fill, 2.6×10^4^ ± 1×10^4^ CFUs, *Shigella* – green fill, 2.6×10^4^ ± 1×10^4^ CFUs) at 6 hp2i. Data pooled from 3 experiments with n>10 larvae per condition per experiment (mean ± SD, ****p < 0.0001, 2-way ANOVA with Sidak’s multiple comparisons test).

**F** Log_10_-transformed CFU counts of naïve and *Shigella*-primed (1.4×10^3^ ± 6.7×10^2^ CFUs) larvae injected with a lethal dose of *S. flexneri* M90T (PBS – black fill, 2.6×10^4^ ± 1×10^4^ CFUs, Shigella – green fill, 2.6×10^4^ ± 1×10^4^ CFUs) and treated with DPI. Data pooled from 3 independent experiments using n=3 larvae per condition per experiment (mean ± SD, 2-way ANOVA with Sidak’s multiple comparisons test).

**G** Normalised fluorescence of GFP driven from *mpx* expression in neutrophils from naïve and *Shigella*-primed larvae (Tg(*mpx*::GFP)) infected with a lethal dose of *S. flexneri* M90T at 6 hp2i. Data pooled from 5 experiments with n>10 larvae per condition per experiment (mean ± SD, ****p < 0.0001, 2-way ANOVA with Sidak’s multiple comparisons test).

